# Concerted Transport and Phosphorylation of Diacylglycerol at ER-PM Contacts Sites Regulates Phospholipid Dynamics During Stress

**DOI:** 10.1101/2024.01.16.575708

**Authors:** Selene García-Hernández, Richard Haslam, José Moya-Cuevas, Rafael Catalá, Louise Michaelson, Vitor Amorim-Silva, Vedrana Marković, Julio Salinas, Johnathan Napier, Yvon Jaillais, Noemí Ruiz-Lopez, Miguel A. Botella

## Abstract

A universal response of plants to environmental stresses is the activation of plasma membrane (PM) phospholipase C (PLC) that hydrolyzes phosphatidylinositol phosphate (PIP) to produce soluble inositol phosphate (IP) and diacylglycerol (DAG). DAG produced in this way can be either phosphorylated by PM diacylglycerol kinases (DGKs) to produce the second messenger phosphatidic acid (PA) or transferred to the endoplasmic reticulum (ER) by the Synaptotagmin 1 (SYT1) protein at ER-PM Contact Sites (CS). In Arabidopsis, the clearance of DAG at the PM (avoiding deleterious accumulation) by the transfer activity of SYT is essential to maintain PM stability after stress. In this study we identify that DGK1 and DGK2 form a module with SYT1 at ER-PM CS through interaction of their C1 and C2 domains respectively. Global transcriptomic and metabolomic analyses confirms that *SYT1* and *DGK1*/*DGK2* are functionally related and lipidomic analysis supports the hypothesis that DGK1 and DGK2 function at the ER by phosphorylating DAG transferred by SYT1 from the PM. DGK1 and DGK2 show structural similarity to human DGKε, the DGK isoform that function at ER-PM CS in the phosphoinositide (PI) cycle. Our data indicate that components of the PI cycle are conserved between animals and plants and provide a novel mechanism leading to an increase in the efficiency of the PI cycle by channeling the transport and hydrolysis of DAG at the ER-PM CS.

## INTRODUCTION

Membrane contact sites (MCS) are specialized membrane domains where the membranes of two organelles are at very close apposition without fusion (Fernández-Busnadiego et al., 2015). This short distance, usually 10–30 nm, is possible due to proteins acting as molecular tethers between the lipid bilayers (Eisenberg-Bord et al., 2021; Pérez-Sancho et al., 2016). MCS are a distinctive feature of all eukaryotic cells and are present in almost every organelle, enabling non-vesicular exchange of lipids, small molecules, and signals and fostering coordinated organelle responses to dynamic environments (Chang et al., 2013; Phillips, M. J., & Voeltz, 2016; Prinz, 2014). Most MCS involve contacts of the endoplasmic reticulum (ER) with other organelles including the plasma membrane (PM) known as ER–PM contact sites (ER–PM CS) (Bayer et al., 2017; Scorrano et al., 2019; Wu et al., 2018).

ER–PM CS play a crucial role in the inter-organelle communication in plants, mammals, and yeasts. They allow non-vesicular lipid transport (Sawaki et al., 2016), regulate calcium homeostasis (Saheki & Camilli, 2017), and contribute to the maintenance of cortical ER morphology (Siao et al., 2016). Among the proteins involved in tethering at the ER–PM CS in plants are the synaptotagmins (SYTs) and their counterparts in yeast and mammals known as Extended SYTs (E-Syts) and tricalbins (Tcbs). The Arabidopsis SYT proteins are characterized by a transmembrane (TM) domain at the N-terminus that is embedded in the ER membrane, a SYT-like mitochondrial-lipid binding protein domain (SMP), which can harbor lipids in a hydrophobic cavity (Kopec et al., 2010; Ruiz-Lopez et al., 2021; Saheki et al., 2016; Schauder et al., 2014), and two C2 domains at the C-terminus that allow tethering with negatively charged phosphatidylinositol phosphate (PIP) at the PM (Manford et al., 2012; Pérez-Sancho et al., 2015; Ruiz-Lopez et al., 2021; A. L. Schapire et al., 2008).

Environmental stresses trigger the phospholipase C (PLC)-catalyzed hydrolysis of PIP into soluble inositol phosphates and diacylglycerols (DAG) at the PM (Arisz et al., 2013; Kanehara et al., 2015). Accumulation of DAG may lead to membrane destabilization inducing negative curvatures or even producing membrane fission and fusion dynamics due to the conical structure of these molecules (Campomanes et al., 2019). Therefore, DAG is maintained at a very low concentration (Gaude et al., 2007) in PM but the precise mechanisms for this regulation are still unknown. It is known, according to previous studies, that diacylglycerol kinases (DGK) convert DAG into phosphatidic acid (PA), playing a double role: maintaining DAG levels and playing a role in signaling through the generation of PA (Pokotylo et al., 2018). Recently, it has been described that Arabidopsis SYT1 and SYT3 are involved in the transfer of DAG from PM to the ER (Ruiz-Lopez et al., 2021), opening the possibility that DAG may be phosphorylated into PA in the ER membrane.

In the *Arabidopsis thaliana* genome, there are seven genes encoding *DGKs* (*DGK1-DGK7*) that are divided into three clusters (Foka et al., 2020). DGKs from Clusters I and II (DGK3-DGK7) comprises a catalytic domain with an ATP-binding site responsible for the phosphorylation of DAG (Gómez-Merino et al., 2004), and an accessory domain whose function remains elusive. However, they lack additional domains and therefore are predicted to be cytoplasmic. DGK1 and DGK2 which from Cluster I are unique since they possess a transmembrane region that anchors them to the ER (Vaultier et al., 2008) and two C1 domains mediating DAG binding (Colón-González & Kazanietz, 2006) and protein-protein interactions (Anilkumar et al., 2003). In contrast to plant, human DGKs exhibit a large variety of domains that regulate their activity and membrane targeting (Sakane et al., 2007; S. Xie et al., 2015). Notably, human type III HsDGKε is only anchored to the ER by a TM (Kobayashi et al., 2007), as occurs in DGK1 and DGK2. HsDGKε is an enzyme that catalyzes one of the steps of the phosphatidylinositol (PI) cycle, the major metabolic pathway for the synthesis of phosphatidylinositol (PI) and its phosphorylated forms (Bozelli & Epand, 2019). Therefore, this cycle is essential to replenish the PIP used by PLC at the PM, and to sustain repetitive rounds in response to physiological stimuli. Interestingly, this metabolic cycle is atypical as it occurs in two different cellular compartments i.e., the ER and the PM, implying that some lipids species must be transferred between the ER and the PM, and vice versa. This characteristic enables the ER-PM CS to have an essential role in this process (Chen et al., 2019; Lin et al., 2022).

In this work, using protein-protein interaction studies, confocal microscopy, transcriptomics, metabolomics, lipidomics, and phenotypic characterization, we describe the mechanism by which DAG produced by stress-activated PLC is removed from the PM by the combined action of SYT1, DGK1, and DGK2. We report that the DAG transferred from the PM to the ER by SYT1 is used by DGK1 and DGK2 channeled by the interaction of SYT1 and DGK at ER-PM CS. We speculate that this process is part of the PI cycle, a sequence of enzymatic reactions that is central to many cellular functions, including cell growth, stress resistance, cytoskeletal organization, and vesicular trafficking (Chang et al., 2013, 2017).

## RESULTS

### SYT1 interacts with DGK1 and DGK2 at ER-PM CS

SYT1 is an ER-PM tether that plays a role in maintaining diacylglycerol (DAG) homeostasis at the plasma membrane (Ruiz-Lopez et al., 2021). As a result of a yeast two-hybrid screening using the C2 domains of SYT1 as bait (amino acids 244-541), we identified a clone corresponding to the amino acids 79-515 of the Diacylglycerol Kinase 2 (DGK2) (AT5G63770), a protein involved in the conversion of DAG to phosphatidic acid (PA). In addition, we independently identified the highly related DGK1 (AT5G07920) in via TAP-TAG screening using SYT1 as a bait. Thus, we decided to investigate this putative interaction further, as SYT1 and DGKs share a function related to DAG. Using *Nicotiana benthamiana* we found that a full-length SYT1 protein interacted with DGK2 in both Co-IP (Fig. 1A) and BiFC assays indicating that this interaction takes place at ER-PM CS (Fig. S1). We also noticed that SYT1 associate with DGK1 using Co-IP (Fig. 1B). Moreover, our results showed that DGK1 immunoprecipitated DGK2 *in vivo*, supporting the hypothesis that SYT1, DGK1, and DGK2 likely form a complex at the ER-PM CS (Fig. 1A, Fig. 1B, Fig. 1C).

**Figure 1.**
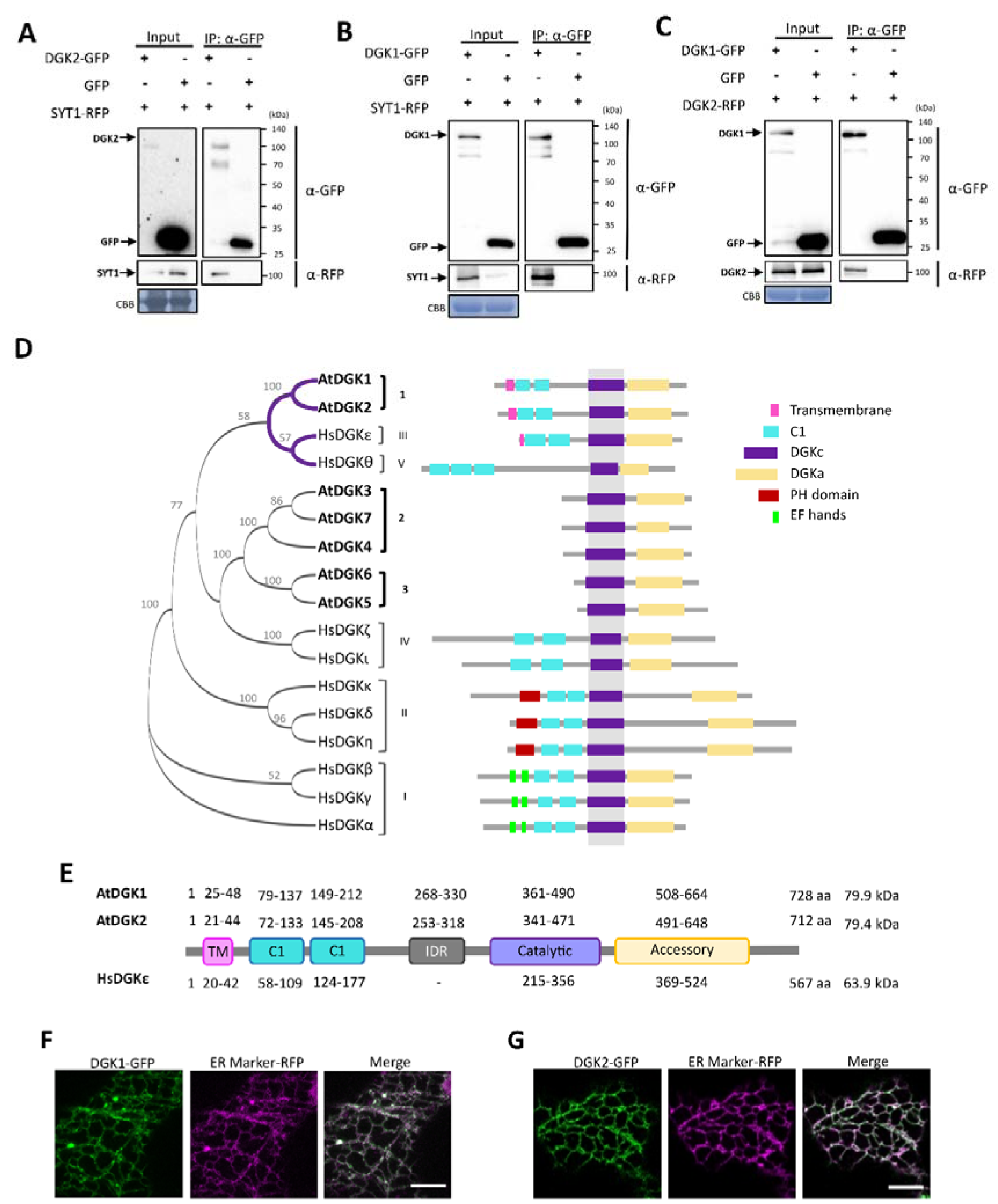
DGK1 and DGK2 (Cluster I) interact with SYT1, and both share a similar structure and ER localization. **A-C)** A GFP pull-down assay shows that SYT1, DGK1, and DGK2 are part of the same complex in which SYT1 interacts with DGK2 (A) and with DGK1 (B). Additionally, DGK1 and DGK2 interact with each other (C). Proteins were transiently coexpressed in *N. benthamiana*, and tissue was harvested 2 days post-infiltration. Proteins tagged with GFP were immunoprecipitated using GFP Trap beads. Total (input) and immunoprecipitated (IP) proteins were separated by SDS-PAGE, and DGK2-GFP andDGK1-GFP were detected with a GFP antibody. SYT1-RFP and DGK2-RFP were detected using an anti-RFP antibody. Uniform sample loading was verified by Coomassie blue staining (CBB) on the input samples. **D)** Phylogenetic tree of proteins from the DGK families of Arabidopsis and Humans, along with the protein structures featuring schematized domains. The evolutionary history was inferred using the Maximum Likelihood method and the JTT matrix-based model. The percentage of replicate trees in which the associated taxa clustered together in the bootstrap test is displayed adjacent to the branches. **E)** Schematic view of DGK1, DGK2, and HsDGKε proteins with domain data extracted from the UniProt database. Domain boundaries are indicated by the corresponding amino acid numbers. Notably, DGK1 and DGK2 feature an IDR, while HsDGKε does not. **F)** DGK1, DGK2, and the ER marker (FaFAH) were tagged with GFP or RFP at the C-terminus and expressed in *N. benthamiana* leaves. Both DGK1 and DGK2 exhibit a reticular pattern due to their anchoring to the ER via the TM domain. Confocal images were captured in the cortical region of lower epidermis cells at 2 days post-infiltration. Scale bar = 10 µm.

The Arabidopsis DGK family comprises seven genes (DGK1-7) that cluster into three clades in a phylogenetic analysis (Fig. 1D). Cluster I is composed by the large proteins DGK1 and DGK2, Cluster II contains DGK3, DGK4, and DGK7, and Cluster III encompasses the closely related DGK5 and DGK6 (Gómez-Merino et al., 2017). Expression of Arabidopsis *DGK* genes in vegetative tissues using RNA sequencing data from eFP SeqBrowser indicated that *DGK1*, *DGK2*, *DGK3*, *DGK5* and *DGK7* are ubiquitous, whereas *DGK4* and *DGK6* are mainly expressed in pollen (Fig. S2A, Fig. S2B). A phylogenetic analysis of the identified domains of the Arabidopsis and the ten human DGKs is shown in Fig. 1D. All DGKs possess a catalytic and an accessory domain with unknown function. Notably, only DGK1 and DGK2 are closely related to human clusters III and IV, which include HsDGKε and HsDGKθ. Arabidopsis clusters II and III are most related to clusters III and IV in humans that encompass HsDGKζ and HsDGKι (Fig. 1D). All human DGKs possess two (or three in the case of HsDGKθ) C1 domains that bind DAG and phorbol ester. In contrast, these domains are exclusive to the Arabidopsis DGK1 and DGK2 (Fig. 1D). Interestingly, only the Arabidopsis DGK1, DGK2, and the closely related HsDGKε contain a transmembrane domain (TM) that target these proteins to the endoplasmic reticulum (Kobayashi et al., 2007, Vaultier et al., 2008), as consequence. Arabidopsis DGKs in clusters II and III are predicted to be cytosolic. Therefore, these proteins should be recruited at particular cellular locations to act on specific DAG pools, such as DGK4 that is localized at the ER (Angkawijaya et al., 2020), even though its sequence does not contain a predicted TM (Fig. 1D). A more detailed analysis of DGK1, DGK2, and HsDGKε revealed that they possess a TM at the N-terminus, two C1 domains, a catalytic domain, and an accessory domain at the C-terminus in similar positions (Fig. 1E). The main difference between Arabidopsis DGK1 and DGK2 with HsDGKε lies in the presence of an Intrinsic Disordered Region (IDR) in DGK1 and DGK2 between the C1 and the catalytic domains that is not predicted in HsDGKε (Fig 1E, Fig. S3A, Fig. S3B). DGK1 and DGK2 localized at the ER, localization that is depended on their TM (Angkawijaya et al., 2020; Vaultier et al., 2008). Consistently, transient co-expression of DGK1-GFP and DGK2-GFP with a bulk ER-marker in *N. benthamiana* showed complete colocalization, indicating that DGK1 and DGK2 proteins localize throughout the ER when ectopically expressed (Fig. 1F).

The interaction observed between DGK1 and DGK2 with SYT1 hints a function of these kinases at ER-PM CS, a localization recently reported for the human homolog HsDGKε (Bozelli & Epand, 2019; Hozumi et al., 2017; Tao-Cheng, 2018). To investigate this further, we transiently co-expressed DGK1 together with DGK2 and SYT1 in *N. benthamiana* leaves. DGK1 and DGK2 showed a bulk ER localization when co-expressed (Fig. 2A, Fig. 2E). However, when DGK1 was co-expressed with SYT1, we found an enrichment of DGK1 at the ER-PM CS (Fig. 2B, Fig. 2E). A similar localization was found for DGK2 when co-expressed with SYT1 (Fig. S4A). SYT1 localization at the ER-PM CS is dependent on the insertion of its TM into the ER and the binding of the C2 domains to negatively charged phospholipids at the PM (Pérez-Sancho et al., 2015, Benavente). Consistently, SYT1 constructs with the SMP domain but lacking the C2 domains (SYT1ΔC2AB) localized in bulk ER when expressed in *N. benthamiana*, and its co-expression with DGK1 did not result in its relocalization at ER-PM CS (Fig. 2C). Similarly, we did not observe ER-PM CS localization of an ER marker when co-expressed with SYT1 (Fig. S4B). Furthermore, co-expression of the artificial ER-PM Contact Sites MAPPER (Lee et al., 2019) with DGK1 or DGK2 did not induce the localization of these proteins at ER-PM CS (Fig. 2D, Fig S4C). Collectively these experiments indicate that the enrichment of DGK at the ER-PM CS depends on the interaction with SYT1 and is not caused by a putative increase in the formation of ER-PM CS.

**Figure 2.**
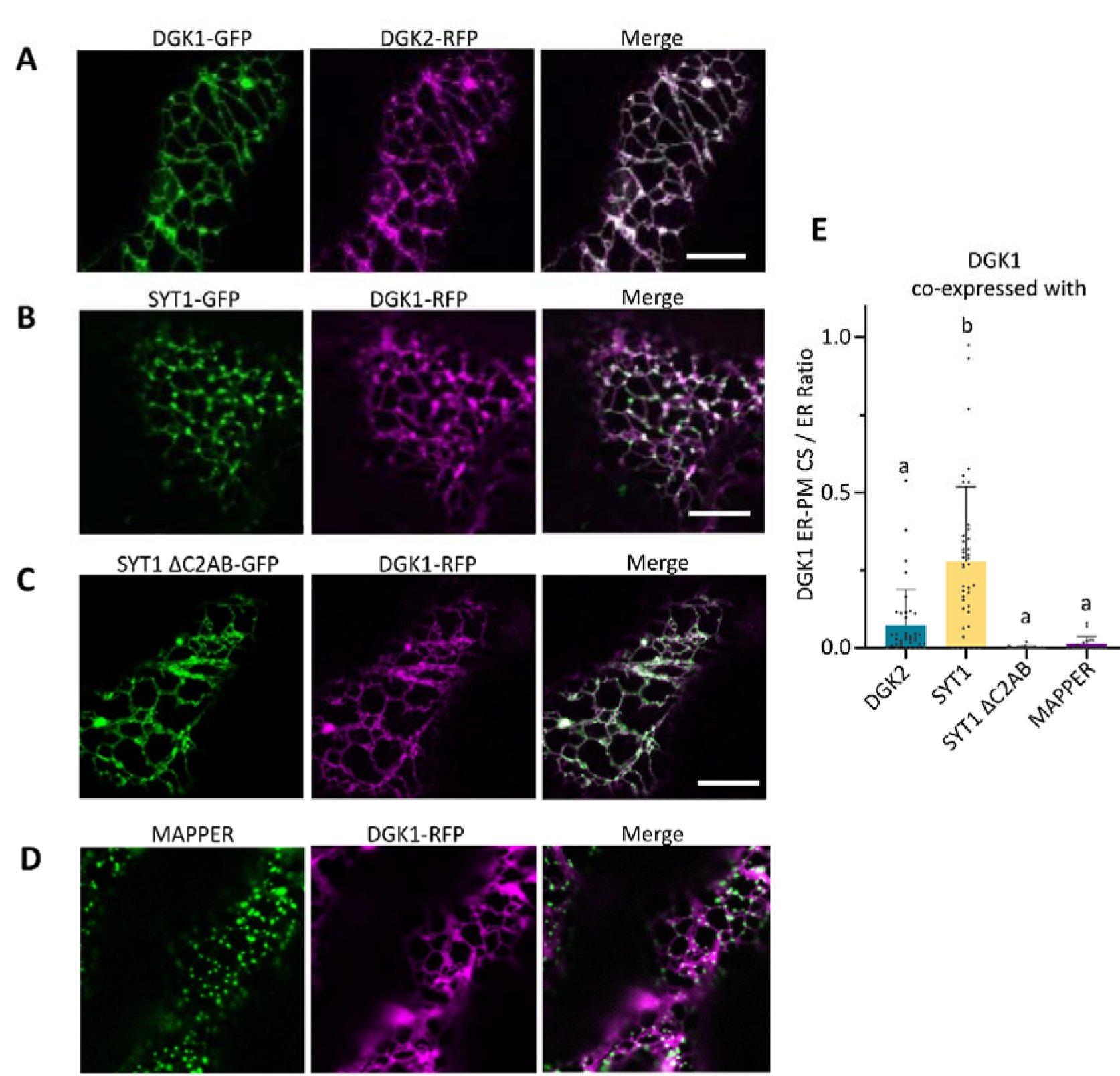
SYT1 drags DGK1 to the ER-PM CS. **A to D)** DGK1 was co-expressed with DGK2 (A), SYT1 (B), SYT1 ΔC2AB (C), and MAPPER (D), in *N. benthamiana* leaves. The cortical region of lower epidermal cells from 2 days-post infiltration leaves is shown as individual channels of each protein and merged. Scale bar = 10 µm. **E)** The DGK1 ER-PM CS / ER ratio was quantified in plants co-expressing DGK1 with DGK2, SYT1, SYT1 ΔC2AB, or MAPPER. When DGK1 was co-expressed with SYT1, DGK1 increased at the ER-PM CS but not when was co-expressed with SYT1 ΔC2AB or MAPPER. ER-PM CS and ER segmentation were generated using machine learning (Ilastik). Letters indicate statistically significant differences using one-way ANOVA Tukey multiple comparisons, p < 0.05 (n > 20).

### Identification of the interaction domains of DGK1/DGK2 and SYT1

To identify the domains of DGK1 required for interacting with SYT1, we generated truncated forms of DGK1 and SYT1 and conducted Förster Resonance Energy Transfer (FRET) experiments (Fig 3A). While full length SYT1-GFP localized at ER-PM CS, SYT1 lacking the C2 domains (SYT1ΔC2AB-GFP) and incapable of PM binding, localized at bulk ER (Fig 3C). Full-length DGK1-GFP, DGK1 without the accessory domain (DGK1ΔAcc-GFP), and DGK1 containing the TM and the C1 domains (DGK1-TM-C1- GFP) showed bulk ER localization (Fig 3B). FRET analysis confirmed the *in vivo* interaction between SYT1 and DGK1 (both C-terminal tagged to GFP or RFP) when co-expressed in *N. benthamiana* leaves (Fig 3D). Conversely, and consistent with the Y2H data, the SYT1 protein lacking the C2 domains (SYT1ΔC2AB) did not interact with DGK1, despite their colocalization at bulk ER. SYT1 showed a similar interaction with DGK1, DGK1ΔAcc, and DGK1-TM-C1s that in addition to the TM only contain the C1 domains (Fig 3D), implying that the C2 domains of SYT1 interact with the C1 domains of DGK1.

**Figure 3.**
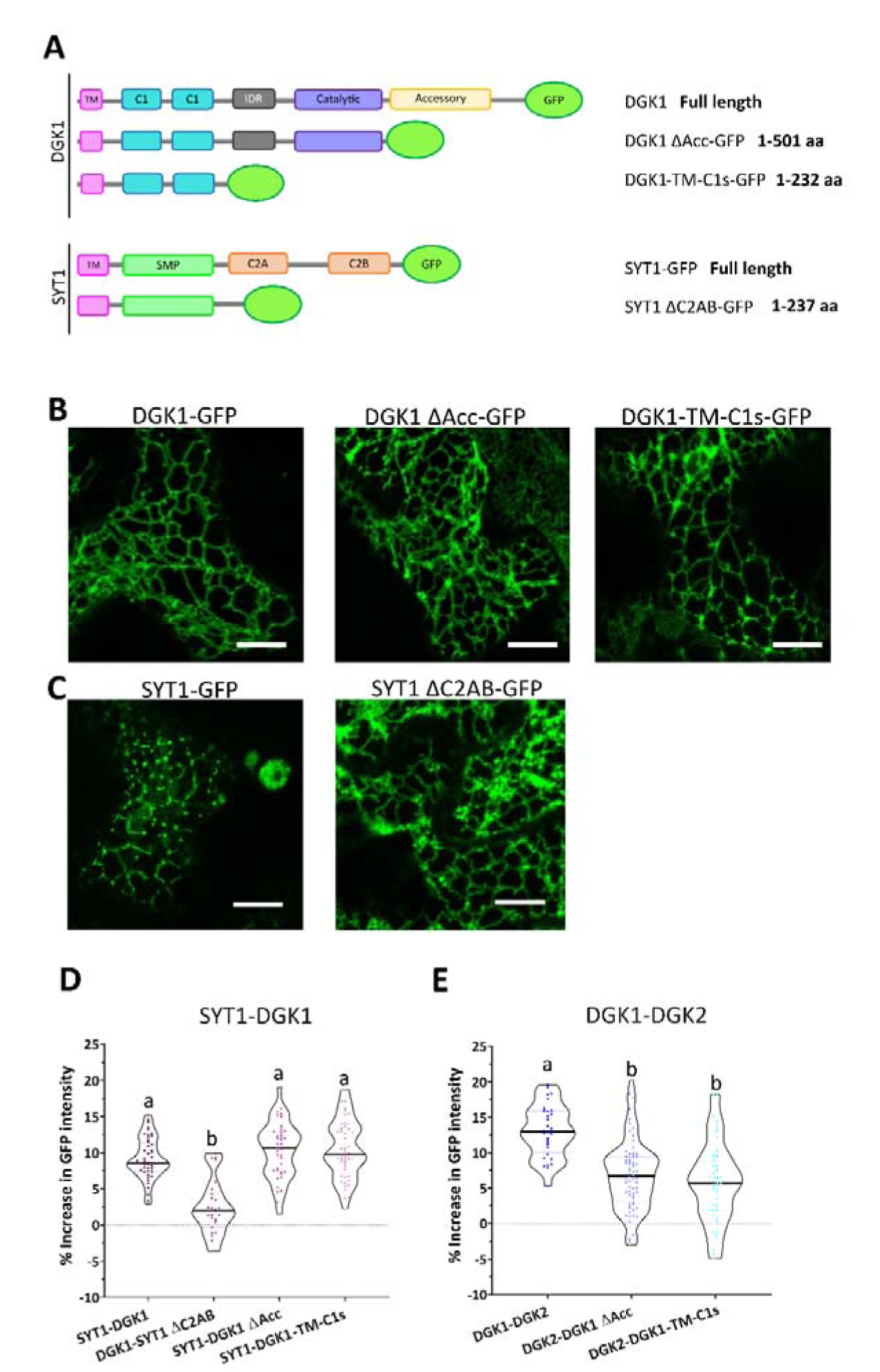
DGK1 and SYT1 interact through C1 and C2 domains whereas DGK1 and DGK2 bind thanks to the Accessory domain. **A)** Schematic view of SYT1, DGK1 and their truncated protein versions (DGK1 ΔAcc, DGK1 TM-C1s and SYT1 ΔC2AB) tagged to GFP at the C-terminus. The first and last amino acids are represented. **B and C)** The localization of both full-length and truncated versions of DGK1 (B) and SYT1 (C) were observed in *N. benthamiana* epidermal cells using confocal microscopy at 2 days post-infiltration. Scale bar = 10 µm. **D and E)** Förster resonance energy transfer (FRET) assays were conducted using complete or truncated versions of SYT1, DGK1, and DGK2 to determine the domains responsible for the interaction between SYT1 and DGK1 (A) and between DGK1 and DGK2 (B). Pairs of proteins were co- expressed in *N. benthamiana* leaves and analyzed at 2 days post-infiltration. RFP-tagged proteins were photobleached, and GFP-tagged proteins were quantified. The percentage increase in GFP intensity was calculated using the following formula: [(IAfter − IBefore) / IAfter] × 100, where IBefore and IAfter represent the means of the intensity from 6 measurements taken before and after photobleaching, respectively. The lowercase letters indicate statistically significant differences using one-way ANOVA and Tukey multiple comparisons (p < 0.05).

To investigate the interacting domains of DGK1 and DGK2, we performed FRET between DGK2 and the DGK1 deletion constructs (Fig 3A). While full-length DGK2 and DGK1 shows interaction, DGK2 did not interact with DGK1ΔAcc (Fig 3E), supporting the conclusion that DGK1 and DGK2 proteins interact through their accessory domains. Because all DGKs we have analyzed so far contained an accessory domain (Fig 1D), it is tempting to speculate that the accessory domains of DGKs may play a role in their localization by the formation of homo- and/or heterodimers.

### Mutations in *DGK1*, *DGK2,* and *SYT1* cause similar changes in their transcriptional and primary metabolite levels

To determine the functional relationship of *DGK1*, *DGK2*, and *SYT1*, the gene regulatory network in their respective mutants was analyzed. Homozygous T-DNA mutants were produced for *DGK1* (*dgk1- 1*, SALK_053412, *dgk1*), *DGK2* (*dgk2-2*, SAIL_71_B03, *dgk2*) (Fig. S5A), and *dgk1dgk2* by crossing *dgk1* and *dgk2* (Fig. S5B). No transcripts were detected by RT-PCR (Fig. S5C), and they are expected to be null mutants because the T-DNAs disrupt the catalytic domains of DGK1 and DGK2 (Fig. S5D).

No clear phenotypic differences were found between the double *dgk1dgk2*, *syt1* (Ruiz-Lopez et al., 2021) and the wild type (WT) under standard growth conditions in any of the tissues or developmental stages analyzed (Fig. S5E, Fig. S5F, and Fig S5G). Nevertheless, RNA-sequencing identified a total of 227 (118 up- and 109 down-regulated) and 217 (183 up- and 34 down-regulated) differentially expressed genes (DEGs) in *dgk1dk2* and *syt1* vs WT, respectively (Fig 4A), listed in Supplemental Table S1. When we compared DEGs between *dgk1dgk2* and *syt1*, we found a non-random overlap (49.7 enrichment with a *p-*value of 4,2e^-106^) of 73 genes, which accounts for 32.16% of those identified in the *dgk1dgk2*. Of the 118 genes induced in *dgk1dgk2,* 66 were also induced in *syt1,* with a remarkable 56% overlap (102.6 enrichment with a *p-*value of 5,7e^-122^) (Fig. 4B). The enrichment was calculated as the number of overlapping genes divided by the expected number of overlapping genes drawn from both groups. The striking similarity of de-regulated genes by *DGK1*/*DGK2* and *SYT1* strongly suggests that *SYT1* and *DGK1*/*DGK2* are functionally related. Functional enrichment analysis of the upregulated genes for *dgk1dgk2* and *syt1* (Supplemental Table S2), unveiled the highest over-representation of GO terms related to jasmonic acid (JA) biosynthesis and signaling, a hormone involved in wound responses (Sheard et al., 2010), with considerable overlap of the top enriched 15 GO terms in all genotypes (Fig 4C, Fig. 4D).

**Figure 4.**
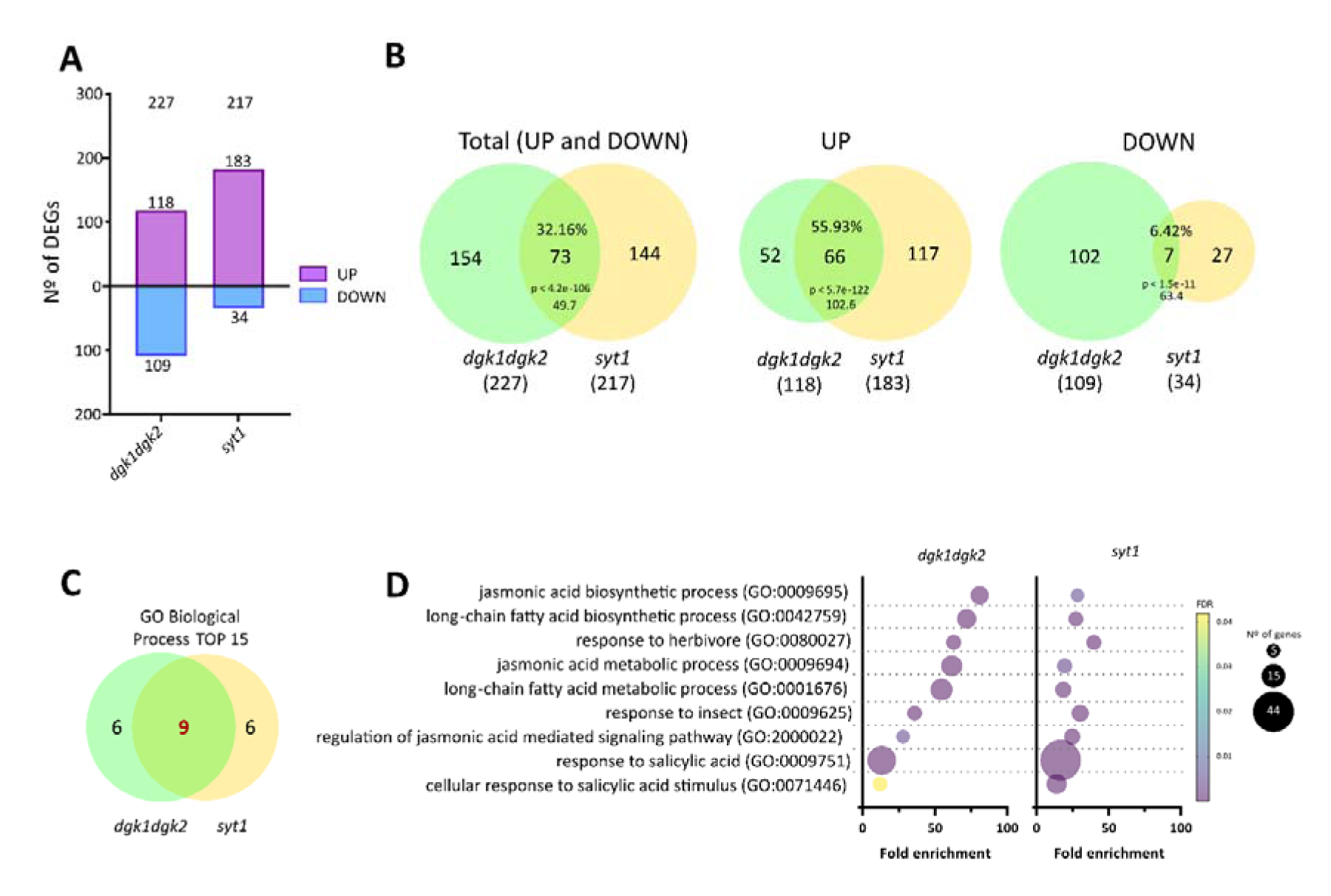
*SYT1* and *DGK1/2* modulate a similar set of genes. RNAseq analysis was performed using the aerial parts of a pool of 2-week-old seedlings (10 seedlings per biological replicate, and at least 3 replicates per group) grown in soil under control condition. **A)** *dgk1dgk2* and *syt1* differentially expressed genes DEGs (q-value < 0.05), compared to the wild type (WT), are displayed. Up-regulated genes are represented in purple, and down-regulated in blue. **B)** Schematic representation of the total, up-regulated, and down-regulated genes in *dgk1dgk2* and *syt1* mutants and its overlap. The percentages of shared genes are calculated relative to the number of total DEGs in the *dgk1dgk2* mutant. The enrichment of the overlap (calculated as the number of overlapping genes divided by the expected number of overlapping genes drawn from two independent groups) is 49.7, 102.6 and 63.4 for Total, up, and down-regulated genes, respectively. The expected genes were defined by next formula: (n° of genes in *dgk1dgk2* * n° of genes in *syt1)* /total genes analyzed in full genome. p-value of the enrichment is indicated. Notably, *DGK1/2* and *SYT1* appear to activate the same pathway, as their mutants share 56% of up-regulated genes. **C)** Venn diagram showing the overlap between the top 15 GO terms associated with up-regulated genes in both genotypes. They share 9 GO terms (in red). **D)** Bubble plot of the 9 GO terms shared in panel C. The fold enrichment of each category is shown on the X axis, while the False Discovery Rate (FDR) is represented by a color gradation from purple, lowest, to yellow, higher. The size of the circle corresponds to the number of genes involved in each GO term. Most of these GO terms are related to Jasmonic Acid pathway.

We also analyzed changes in components of the primary metabolism for the *dgk1dgk2* and *syt1.* A total of 32 primary metabolites were quantified by gas chromatography-mass spectrometry (GC–MS) including amino acids, sugars, sugar derivates, and organic acids (Supplemental Table 3). In the *dgk1dgk2* double mutant, there was an increase of 11 out of the 32 primary metabolites, sharing seven with *syt1* (Fig. 5A), that had 15 significantly changed metabolites compared to WT (14 increased and 1 decreased). Both mutants share the trend of increase or decrease in most metabolites (Fig. 5B), further supporting the related function of DGK1/DGK2 and SYT1.

**Figure 5.**
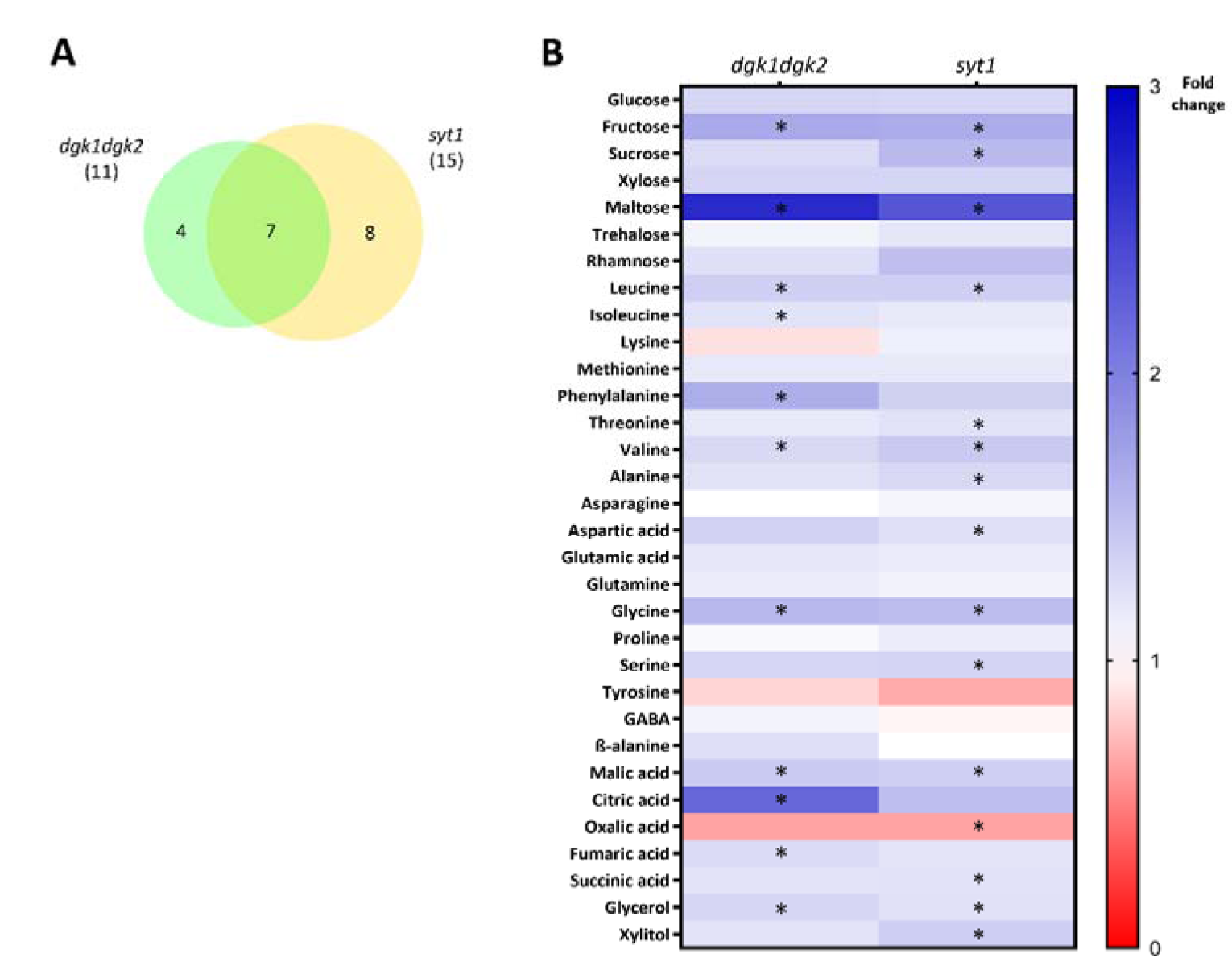
Primary metabolite analysis shows that *dgk1dgk2* and *syt1* have a similar accumulation compared to WT. Primary Metabolomic analysis was performed using the same tissue than in RNAseq: the aerial parts of a pool of 2-week-old seedlings (10 seedlings per biological replicate, and 4 replicates per group) grown in soil under control condition. **A)** Venn diagram represents the primary metabolites significantly different in both mutant from the WT, determined using the multiple t-test (Holm-Sidak) method (p < 0.05). 11 out of the 32 analyzed metabolites are affected in *dgk1dgk2* and 15 in *syt1.* They share 7 of these metabolites. **B)** Heatmap displaying fold change of the metabolic alteration in the mutants compared to WT. Red indicates those metabolites repressed; White the ones that do not change; Blue those that increase. Asterisks indicate the metabolites significantly different from WT summarized in A).

### DGK1 and DGK2 function at the Endoplasmic Reticulum

DGK1 and DGK2 are anchored to the ER and form a complex with SYT1 at ER-PM CS to regulate DAG homeostasis. Therefore, their activity is presumed to occur in *cis*, i.e., in the ER membrane where it is anchored. However, the interaction of DGK1 and DGK2 with SYT1 at ER-PM CS and the presence of an IDR preceding the catalytic domain raise the possibility that DGK1 and DGK2 function in *trans* by catalyzing the phosphorylation of DAG at the PM.

To address whether DGK1 and DGK2 function in *cis* or in *trans*, we isolated membrane fractions enriched in plasma membrane (PM) and inner membrane (IM) from leaves of WT and *dgk1dgk2* plants growing in control conditions and after cold-treatment, by employing a two-phase partitioning protocol (Ruiz-Lopez et al., 2021). We used cold stress because this condition i) enhances the production of DAG by activating phospholipase C (PLC) (Vergnolle et al., 2005), ii) promotes the relocalization of SYT1 and SYT3 at the plasma membrane (Ruiz-Lopez et al., 2021), iii) induced the accumulation of DAG at the PM in the *syt1syt3* double mutant (Ruiz-Lopez et al., 2021), and iv) induced the accumulation of *DGK1* and *DGK2* transcripts (Fig. S2C, Fig. S2D) (Gómez-Merino et al., 2004). To verify the efficiency of the purification method, we analyzed the initial total extract (T), the crude microsomal fraction (M), the enriched PM samples, and the inner membranes (IM) by immunoblots using antibodies of protein markers from different cellular compartments (Fig. S6). The PM fraction showed a pronounced enrichment of a PM marker and a significant reduction of ER, tonoplast, and chloroplast markers.

We then utilized liquid chromatography tandem mass spectrometry (LC–MS; High Resolution / Accurate Mass) on the PM and IM lipid fractions, profiling 63 molecular species spanning three glycerolipid classes, DAG, and the abundant phosphatidylcholine (PC) and phosphatidylethanolamine (PE) (Supplemental Table S4). The analysis did not reveal changes in the PM composition of PC or PE between the WT and *dgk1dgk2* under control conditions, and a slight increase of DAG that was only significant in total DAG (Fig. S7). When analyzed the IM no significant changes in any of the lipids were obtained. In cold-stressed plants, no changes of DAG, PC or PE were found in the PM (Fig. 6). However, a significant increase in DAG levels (1.6 enrichment) and a slight increase in PC and PE (1.2 and 1.1, respectively) was obtained in the IM of the *dgk1dgk2* mutant compared to WT (Fig. 6). Single lipid species analysis revealed that most DAG molecular species at the IM displayed a tendency to accumulate in *dgk1dgk2* compared to WT, being DAG34:2 and DAG34:4 the most enriched species (1.8 enrichment) (Fig. 6). Additionally, PC34:3 (1.4 times) and PE34:2 and PE34:3 (1.2 times) species were also enriched in *dgk1dgk2* compared to WT. As DAG serves as the substrate for PC and PE biosynthesis at the ER through the CDP-choline and CDP-ethanolamine pathways, the increased levels of these two glycerolipid classes in the IM of *dgk1dgk2* plants might be an indirect result of the increased abundance of DAG in these plants. These results strongly support that DGK1 and DGK2 function in *cis* by phosphorylating DAG molecules at the ER.

**Figure 6.**
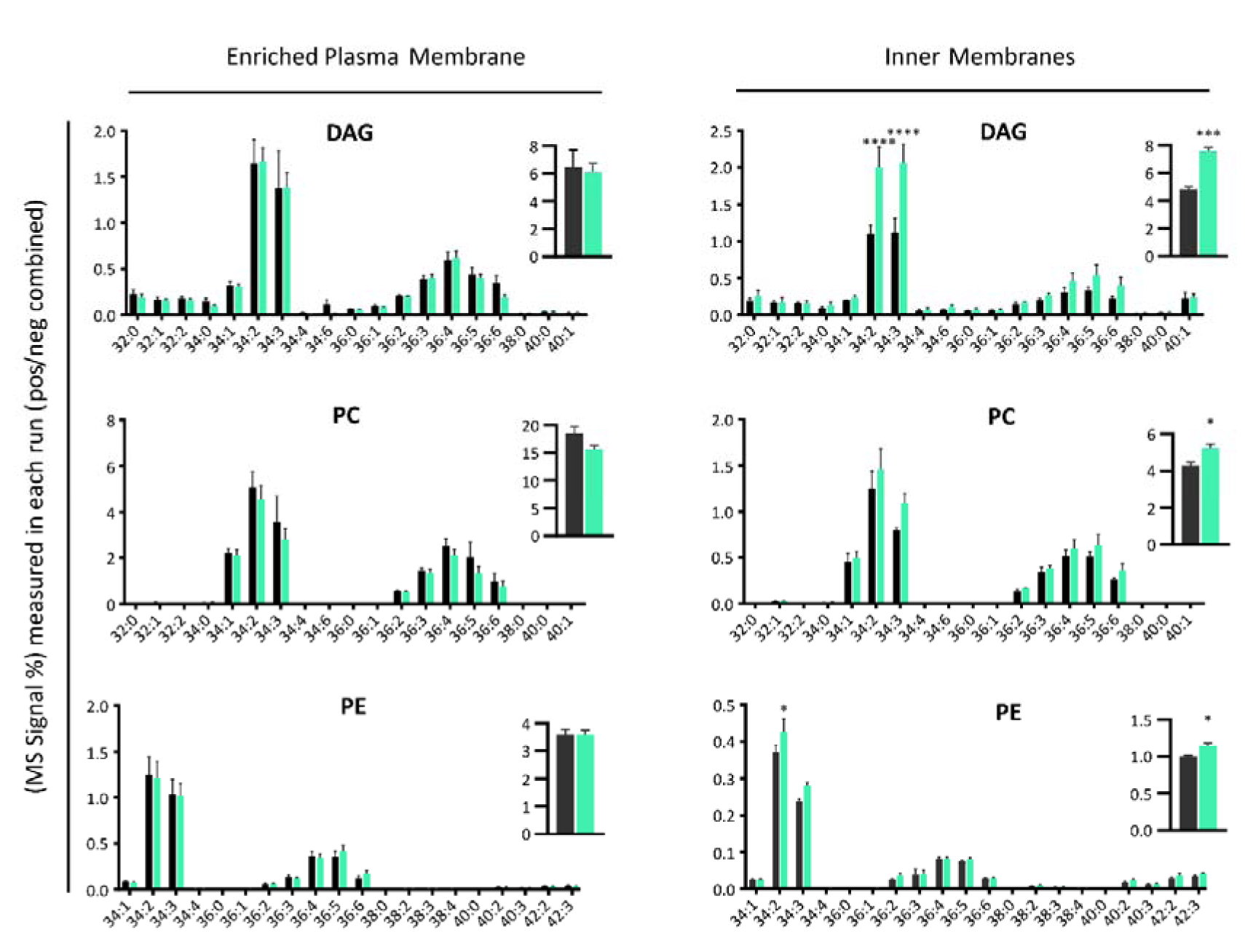
DAG accumulates at the Inner Membranes (IM) in the *dgk1dgk2* mutant. High-resolution/accurate mass (HR/AM) lipid analysis of the molecular species of DAG, PC and PE in Plasma Membrane (PM) and Inner Membranes (IM). Fractions were isolated from 5-week-old WT and *dgk1dgk2* rosettes grown at control condition followed by 3 days of cold treatment (4°C). PM and IM samples were purified by two phase partitioning protocol and lipids were extracted following as described in ‘Methods’. Acyl chains are expressed as number of acyl carbons: number of acyl double bonds. Distribution of the identified DAG, PC, and PE molecular species in the PM and in the IM of WT (gray) and *dgk1dgk2* (green) is represented. Column bars show the mean values of at least three biological replicates, with error bars indicating the standard error of the mean (SEM). The asterisks indicate statistically significant differences between *dgk1dgk2* and WT as determined by a Dunnett’s multiple comparisons test: ****p < 0,0001; ***p < 0,0002; **p < 0,0021; *p < 0,0332.

### The *DGK2* gene plays a role in cold stress root growth and cold-acclimated freezing tolerance

*SYT1* is involved in the resistance to various abiotic stresses (Pérez-Sancho et al., 2015; Arnaldo L. Schapire et al., 2008; Yamazaki et al., 2008). In addition to *DGK1* and *DGK2* genes being induced by cold (Fig. S2C, Fig.S2D), the *dkg1dgk2* mutant showed differences in their lipidome after cold stress. Therefore, we investigated whether *DGK1* and *DGK2* play a role in cold stress resistance.

First, we investigated if the induction of *DGK1* and *DGK2* transcripts corresponded to an increase in the amount of protein. For this, the coding region of *DGK1* and *DGK2* fused to *RFP* and *GFP,* respectively, and were driven by their own promoters. These constructs were then transformed into their corresponding mutants (Fig. S8A, Fig. S8B). Further analysis confirm the presence of the corresponding DGK1-RFP and DGK2-GFP fusion proteins in seedlings, adult leaves, and internodes using immunoblot analyses (Fig. S8C, Fig. S8D). Notably internodes showed the highest content of DGK1-RFP and DGK2-GFP, consistent with their transcriptome data (Fig. S2B). Further analysis into the levels of DGK1-RFP and DGK2-GFP in seedlings, leaves and internodes was conducted under control conditions and after cold treatment. In both lines and across the three tissues analyzed, there was an increase in the amount of protein abundance after cold stress (Fig. 7A, Fig. 7B). The induction of DGK1 and DGK2 proteins was dependent on the tissue, increasing in leaves more than seven-fold for DGK1 and more than two-fold for DGK2 (Fig. 7A, Fig. 7B). Additionally, the consistent appearance of a smaller band in DGK1 after cold treatment suggest that it could be caused by an alternative splice variant.

**Figure 7.**
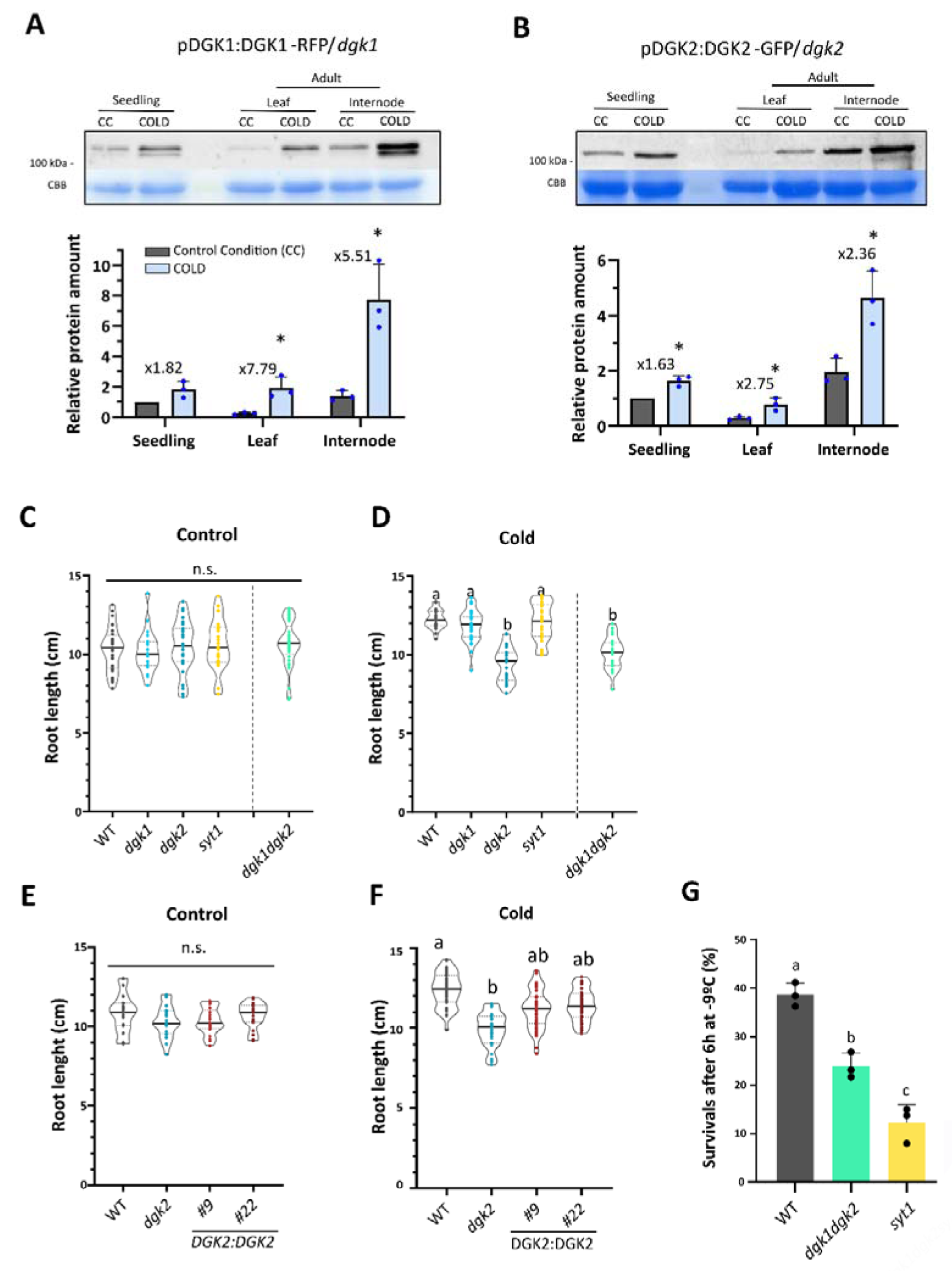
*DGK2* is involved in cold response and plays an additive role with *SYT1* in freezing tolerance. **A and B)** Immunoblot assays show that the protein levels of DGK1:DGK1-RFP (A) and DGK2:DGK2- GFP (B) increase under cold (4°C) treatment in several analyzed tissues, with the internode showing the highest expression and the adult leaf exhibiting the most significant increase after treatment. Measurements were normalized to the seedling control. 2-week-old seedlings were treated for 24 hours, and adult tissues (7 week) were treated for 3 days maintaining the photoperiod. **C and D)** For root elongation assays, seeds of different genotypes were germinated on MS ½ plates at 22°C for 3 days and then transferred to other plates to ensure uniform seedling size. These plates were randomly divided into Control (kept at 22°C for 15 days) (C) and Cold (moved to 10°C for 28 days) conditions (D). While there was no phenotype observed in the control condition (C), in the cold condition (D), *dgk2,* but not *dgk1* nor *syt1*, showed a shorter root length. n > 20. The letters indicate statistically significant differences using one-way ANOVA and Tukey multiple comparisons test (p < 0.05). **E)** Plants directly sown into soil and grown under control conditions for 2-week-old were acclimated for 7 days at 4°C. Subsequently, they underwent a freezing treatment, during which the temperature was decreased by 1°C every 30 minutes until it reached -9°C. This temperature was then maintained for 6 hours before being gradually increased by 1°C every 30 minutes until returning to 4°C. Following this, the plants were transferred back to the control chamber and allowed to recover for 7 days. Afterward, the count of surviving plants was conducted. Column bars represent mean values of three biological replicates (> 20 plants per replicate) and error bars are showing SD. Letters indicate statistically significant differences using one-way ANOVA and Tukey multiple comparisons, p < 0.05.

Despite multiple attempts to visualize the subcellular localization and expression pattern of DGK1- RFP and DGK2-GFP using confocal microscopy, both lines exhibited almost undetectable fluorescence signal in vegetative tissues unabling us to perform further investigation of the subcellular localization. Therefore, to obtain lines with a higher expression that could fully complement the cold phenotype (see below) and be visualized using confocal microscopy, DGK2-GFP lines driven by the CaMV-35S and the UBQ10 promoters were generated. However, although we obtained multiple transgenic lines (Fig. S9A), none of these lines showed detectable expression of *DGK2-GFP* by immunoblot (Fig. S9B, Fig. S9C).

Based on the expression of *DGK1* and *DGK2,* the cold accumulation of DAG in the *dgk1dgk2* mutant, and the root defect shown by plants subjected to the DGKs inhibitor (Gómez-Merino et al., 2005), we investigated their role in root growth under cold stress together with *SYT1*. In control conditions, *dgk1*, *dgk2, syt1* and *dgk1dgk2* seedlings show a WT root growth (Fig. 7C, Fig. S10A). However, in seedlings grown at 10°C, *dgk2* but not *dgk1* nor *syt1,* presented a reduction of around 20-25% of root length compared to WT, indicating that *DGK2* but not *DGK1* nor *SYT1*, play a role in root elongation under cold stress (Fig. 7D, Fig. S10B). Moreover, the *dgk1dgk2* double mutant show a growth defect like *dgk2*, indicating that a mutation in *DGK1* does not enhance cold growth sensitivity of *dgk2* (Fig. 7D, Fig. S10B), despite the strong cold induction of *DGK1* transcripts after cold stress (Fig. S2D). We then used two independent transgenic *DGK2-GFP dgk2* lines and showed that while there is no difference in control conditions (Fig. S7E, Fig. S10C), these lines partially complemented the growth defects of *dgk2* (Fig. 7F, Fig. S10D).

Cold acclimation is an adaptive response by which certain plants increase their freezing tolerance after being exposed for some days to low nonfreezing temperatures (Thomashow, 1999). It has been shown that *SYT1* plays a role in cold-acclimated freezing tolerance (Ruiz-Lopez et al., 2021). Therefore, we determine the role of *SYT1* and *DGK1/DGK2* in cold-acclimated freezing tolerance in adult plants. For these experiments, the survival rate of cold-acclimated (7 days at 4°C) WT, *dgk1dgk2*, and *syt1* plants after exposure to freezing temperatures (-9 °C 6 hours) was measured. As previously shown (Ruiz-Lopez et al., 2021), *syt1* plants exhibited a decreased survival rate than WT plants after freezing (Fig. 7G, Fig. S10E). Similarly, *dgk1dgk2* plants also showed a decrease in freezing tolerance compared to WT plants (Fig. 7G, Fig. S10E). Taken together, our data indicate that both *DGK2* and *SYT1* play a role in plant cold-acclimated freezing tolerance.

## DISCUSSION

### Formation of a SYT1/DGK1/DGK2 complex at ER-PM CS

Abiotic stress causes the activation of PLC leading to the production and transient accumulation of DAG at the PM. However, the stationary amount of DAG at the PM must be maintained at a low level since these molecules exhibit a conical shape within the membrane, due to the small polar head, generating regions of negative curvature, inefficient packing, and subsequent instability (Campomanes et al., 2019). To keep low concentrations at the PM, DAG is either converted to the second messenger PA by DGKs at the PM or transported to the ER by SYT1 and later DAG is phosphorylated to PA by DGK1 and DGK2. Unlike SYT1, that localizes at ER-PM CS, DGK1 and DGK2 localize throughout the ER when they are expressed individually or co-expressed with bulk ER proteins or the artificial tether MAPPER. However, DGK1 and DGK2 increased their ER-PM CS localization when co-expressed with SYT1, establishing that it is the interaction with SYT1 and not an increase of contact sites what is driving DGK1 and DGK2 localization to ER-PM CS. Overexpression in COS-7 cells of human HsDGKε, which is phylogenetically and structurally related to DGK1 and DGK2 (Fig. 1D), caused a bulk ER localization (Bozelli & Epand, 2019; Kobayashi et al., 2007; Matsui et al., 2014). However, in tissues showing a high endogenous expression, HsDGKε localizes at ER-PM contact sites such as the subsurface cisterns of Purkinje cells in rat brains (Tao-Cheng, 2018; Bozelli and Epand, 2019). Like HsDGKε, the endogenous localization of DGK1 and DGK2 might differ from the bulk ER reticulated pattern observed after overexpression. However, we failed to detect any signal above background using confocal microscopy in any of the transgenic lines expressing *DGK1:DGK1-GFP* and *DGK2:DGK2-RFP*, even after cold stress treatment that caused the accumulation of the proteins (Fig. 7A, 7B). We also generated stable Arabidopsis lines of *DGK2* driven by *35S* and *UBQ10* promoters, however, none of the transgenic lines produced protein detected by immunoblot. This lack of expression was previously reported by other groups (Vaultier et al., 2008), suggesting tight regulated expression of these proteins.

Analysis of the interacting domains reveals that the C2 domains of SYT1 interact with the C1 domains of DGKs. While C1 domains of specific proteins such as PLC are primarily involved in DAG binding (Topham and Epand, 2009), they also serve as protein-interacting modules. For example, the C1 domains of DGKζ binds Rac1 (Yakubchyk et al., 2005) and β-arrestins (Nelson et al., 2007). As DGK1 and DGK2 are the only DGKs in Arabidopsis containing C1 domains (Fig. 1D), it is unlikely that any other DGK will interact with SYT1. However, the finding that DGK1 and DGK2 form heterodimers through the accessory domain opens the possibility that other kinases could also be part of the complex. In addition, these findings could have significant implications for general DGK function since accessory domains are present in all DGKs we have analyzed.

The clade that includes Arabidopsis DGK1 and DGK2 and human HsDGKε is unique since these proteins contain a TM domain that anchors them to the ER. The TM domain regulates HsDGKε activity by causing a conformational change that brings the active site closer to the membrane (Bozelli et al., 2022). Interestingly, the activity of HsDGKε is regulated by negative membrane curvature and shows very low activity in flat membranes (Bozelli et al., 2018). Whether DGK1 and DGK2 exhibit similar curvature-dependent activity has yet to be established, but if this was the case, a local accumulation of DAG at SYT1/DGK1/DGK2 ER-PM CS might create a negative curvature, hence activating DGK1/DGK2 activity. It is also tempting to speculate that HsDGKε could form a module with the SYT1 homologs Extended Synaptotagmins to regulate DAG homeostasis in mammals at ER-PM CS as we have shown to occur in plants.

### Functional Relationship of DGK1, DGK2, and SYT1

The interaction between SYT1, DGK1, and DGK2, together with their role in DAG homeostasis, implies an interrelated function. The outcome of the transcriptomic analysis strongly reinforces this interdependency. Remarkably, very few genes were deregulated in either *dgk1dgk2* and *syt1* mutants, yet 56% of the genes induced in *dgk1dgk2* were also induced in *syt1*. This overlap has a *p-value* of 5.7e-122 indicating that mutations in *DGK1DGK2* and *SYT1* mutants produce equivalent molecular outcomes. The Gene Ontology analysis of the DEG revealed the activation of JA biosynthesis and signaling in these mutants (Fig 4A). JA is a plant hormone linked to injury and mechanical damage such as wounding and insect attack (Hickman et al., 2017; Huang et al., 2004; Mcconn et al., 1997). The exogenous application of JA induce an increase in primary metabolites, including amino acids, sugars, and tricarboxylic acids (Hendrawati et al., 2006; Meza et al., 2022), which explain these differences. While there are no phenotypic differences in control conditions, we found that *DGK2* is required for root growth during cold stress. Surprisingly, neither *SYT1* nor *DGK1* mutants show a defect in these conditions. However, we found that both, *SYT1* and *DGK2* play a role in cold-acclimated freezing tolerance, indicating that the phenotypic outcome is complex and probably is dependent on the endogenous localization of SYT1, DGK1, and DGK2 proteins.

### The role of DGK1, DGK2 and SYT1 in the PI cycle

Sustained production of inositol 3-phosphate (InsP3) and DAG during stress responses requires a continuous delivery of PI from its site of synthesis in the ER to the PM to maintain PIP levels. In addition, DAG and PA products generated at the PM must be recycled back to the ER as part of the PI cycle. Since PIPs are produced in the PM, and PI is synthesized in the ER, lipid transfer events between the ER and PM must occur, establishing the ER-PM CS as a fundamental part of this process.

Understanding the cellular membrane where DGKs exert their function is critical for comprehending the DAG pool used by these enzymes and their role in DAG metabolism. The interaction of DGK1 and DGK2 with SYT1, their localization at ER-PM CS, and the presence of an IDR would allow their catalytic domains to reach the PM, opening the possibility of function at the PM. However, the lipidomic analysis of *dgk1dgk2* indicates a cold-stress accumulation of DAG at the ER, supporting a cis-function of DGK1 and DGK2, as previously suggested for DGK2 and DGK4 (Angkawijaya et al., 2020). This supports a dynamic interplay between DGK1, DGK2, and SYT1 where the DAG produced at the PM upon activation of PLC is transported to the ER by SYT1 at ER-PM CS and later transformed into PA by action of DGK1 and DGK2 at the ER. This is further supported by the fact that two of the most abundant DAG species (32:2, 34:2) that accumulated at the PM of the *syt1syt3* mutant (Ruiz-Lopez et al., 2021) are also the primary species accumulated in *dgk1dgk2* ER membranes. And this is further reinforced by the finding that the plant SYT ortholog E-Syt1 accumulates at ER-PM CS following G-protein coupled receptor mediated activation of PLC (Kraub & Haucke, 2016; Saheki et al., 2016; B. Xie et al., 2019) and the identification of HsDGKε at ER-PM CS as the enzyme catalyzing one of the steps in the PI cycle at ER-PM CS (Bozelli et al., 2022). We envisage that this sequential function in the PI cycle can be conserved between the E-Syts and HsDGKε in mammals.

In plants, there is no information on the subcellular localization of CDP-DAG and PI synthases involved in PI formation, although this PI is likely transferred from the ER to the PM by a lipid transfer protein. In animals, PI is transferred by TMEM24, an SMP-containing protein localized at ER-PM CS (Lees et al., 2017; Sun et al., 2019; B. Xie et al., 2022). PI is also transported from the ER to the PM by Nir2, which is also required for the transfer of PA from the PM to the ER (Chang et al., 2013; Kim et al., 2016).

In summary, this study reveals DGK1 and DGK2 collaboration with SYT1 at ER-PM CS. Both DGKs operate in the ER, phosphorylating DAG transferred by SYT1 from the PM. Structural similarity to human DGKε suggests conservation in the PI cycle, unveiling a novel mechanism to enhance its efficiency. Future challenges will be the identification of proteins in plants with functions similar to those described in animal and determining if there are other enzymes involved in the PI cycle at ER-PM CS.

## MATERIAL AND METHODS

### Plant Material

In this study, we utilized *Arabidopsis thaliana* (Col-0 ecotype) and *Nicotiana benthamiana* as our experimental materials. Arabidopsis mutant lines used in this study are: *syt1* (AT2G20990) SAIL_775_A08, previously characterized in Pérez-Sancho et al., 2015; *dgk1-1* (AT5G07920) SALK_053412, and *dgk2-2* (AT5G63770) SAIL_71_B03, both obtained from The European Arabidopsis Stock Center (NASC: http://arabidopsis.info/). The presence of the T-DNA insert in each mutant was confirmed using diagnostic PCR, employing allele-specific primers as listed in Supplemental Table S5. This procedure ensured the identification of homozygous plants carrying the insertion. Double mutant *dgk1dgk2* was generated by crossing the single mutants *dgk1-1* and *dgk2-2*.

To generate transgenic lines 35S:DGK2-GFP, UBQ10:DGK2-RFP, DGK2:DGK2-GFP lines 9 and 22 (all in *dgk2-2* background) and DGK1:DGK1-RFP line 12 (in *dgk1-1* background), the methodology described in the section titled "Generation of Arabidopsis transgenic lines" was employed. All Arabidopsis lines are summarized in Supplemental Table S6.

### Plant manipulation and growth conditions

Standard procedures and conditions were employed for Arabidopsis cultivation. Seeds were surface sterilized using chlorine vapors. A mixture of 100 ml of commercial bleach and 3 ml of 37% [w/w] HCl was placed alongside the seeds in a tightly sealed container for 4 hours. Subsequently, the seeds were air-cleared for a minimum of 2 hours in a laminar flow cabinet and then plated under sterile conditions on half-strength Murashige–Skoog medium, supplemented with 1.5% (w/v) sucrose and 0.8% (w/v) physiological agar as solidifying agent.

The plated seeds underwent a 3-day vernalization period at 4°C in darkness. Following this, the plates were transitioned to grow vertically in a chamber with a long-day photoperiod (16 hours light / 8 hours darkness), a photon flux of 130 ± 30 mmol photons m^−2^ s^−1^ and a temperature of 22 ± 1°C unless otherwise stated. When required, seedlings were collected for analysis or, after 7 d of in vitro growth, transferred to soil in pots with organic substrate and vermiculite [4:1 (v/v)] and kept under control condition (130 ± 30 mmol photons m^−2^ s^−1^ and a temperature of 22 ± 1°C). In soil, plants were watered every 2 days. In case of seed collection, they were dried and stored under low-humidity conditions and freshly harvested seeds were used for phenotypic analysis.

### Plasmid constructs

Genomic DNA and cDNA were obtained from Arabidopsis Col-0 and used as templates for amplifying the promoter and Coding DNA Sequence (CDS), respectively, of the target genes. High-fidelity DNA Polymerase (iProof from BioRad #1725301) and primers listed in the Supplemental Table S5 were employed.

Through Multisite Gateway cloning, the promoter regions upstream of the start codon of *DGK1* (2472 bp) and *DGK2* (1438 bp) were inserted via BP reactions (Invitrogen) into a pDONR R4-L1 vector. These, along with pENTR L4-proCaMV35S-R1 (Karimi et al., 2007) and pENTR L4-UBQ10-R1 (Alassimone et al., 2016), constituted the pENTR R4-promoter-L1 constructs. Similarly, the CDS without stop codons of *DGK1* (2184 bp) and *DGK2* (2136 bp) were cloned into the pDONR L1-L2 vector, generating pENTR L1-CDS-L2 constructs. The same process was applied to truncated versions of DGK1 (DGK1ΔAcc, residues 1-501, and DGK1-TM-C1, residues 1-232), and SYT1 (SYT1ΔC2AB, residues 1-237). All pENTR constructs were confirmed by diagnostic PCR, restriction analysis, and sequencing.

Using LR reactions (Invitrogen), the plasmids pENTR L4-promoter-R1, pENTR L1-CDS-L2, and pENTR R2-tag STOP codon-L3 (GFP) (Karimi et al., 2007), and pEDO006 (RFP) were combined with their corresponding pDestination (pDEST) vectors (pGWB5; pH7m34GW; pDEST GW-VYCE and pDEST GW-VYNE (Gehl et al., 2009)) to create the expression plasmids employed in this study (Supplemental Table S7). The resulting combinations were as follows: DGK1:DGK1-RFP and DGK2:DGK2-GFP for stable transformation of Arabidopsis; UBQ10:DGK2-RFP and 35S:DGK2-GFP for both stable transformation of Arabidopsis and transient expression in *N. benthamiana* ; UBQ10:DGK1-RFP, 35S:DGK1-GFP, UBQ10:DGK1ΔAcc-GFP and RFP, UBQ10:DGK1-TM-C1-GFP and RFP, UBQ10:SYT1ΔC2AB-GFP and RFP, 35S:SYT1-VYCE,35S:DGK2-VYCE, and 35S:DGK2-VYNE for transient expression in *Nicotiana benthamiana*.

The constructs 35S:SYT1-GFP, UBQ10:SYT1-RFP, 35S:GFP, and SYT1:MAPPER have been previously described in (Ruiz-Lopez et al., 2021). The pENTR vector containing the CDS of FaFAH1 (Sánchez-Sevilla et al., 2014) was provided by Iraida Amaya and inserted by LR in the pDEST pGWB5 to form 35S:FaFAH1-GFP.

### Generation of Arabidopsis transgenic lines

Expression vector from “Plasmid constructs” section were introduced into *Agrobacterium tumefaciens* strain GV3101::pMP90 through electroporation. The pGWB5 harboring 35S:DGK2-GFP, and the pH7m34GW,0 containing UBQ10:DGK2-RFP, DGK2:DGK2-GFP or DGK1:DGK1-RFP were transformed into Arabidopsis plants by floral dip generating stable transgenic plants. 35S:DGK2-GFP, UBQ10:DGK2-RFP and DGK2:DGK2-GFP were transform into *dgk2-2* mutants. DGK1:DGK1-RFP was transformed into WT plants. WT, instead of *dgk1-1*, was transformed due to delays in obtaining the *dgk1-1* mutant. A pool of two-week-old seedlings from 18 lines of UBQ10:DGK2-RFP and 22 lines of 35S:DGK2-GFP were analyzed through immunodetection, but no detectable signal was found. From the 19 lines of DGK1:DGK1-RFP and 28 lines of DGK2:DGK2-GFP analyzed, DGK1:DGK1-RFP L12 and DGK2:DGK2-GFP L9 and L22 were selected because they had a detectable signal and a single insertion. The DGK1:DGK1-RFP L12 was crossed with the *dgk1-1* mutant once it was obtained. T-DNA and insertion were achieved in homozygosity through diagnostic PCR and hygromycin resistance selection, respectively. The resulting transgenic lines, DGK1:DGK1-RFP L12 *dgk1-1*, DGK2:DGK2-GFP L9 *dgk2-2*, and DGK2:DGK2-GFP L22 *dgk2-2* were employed in this study.

### Arabidopsis eFP Browser Data Analysis

Expression levels of the 7 DGK genes present in Arabidopsis, across various tissues and stages, were extracted from the available RNA-seq data in eFP-Seq Browser website (https://bar.utoronto.ca/eFP-Seq_Browser/) (Sullivan et al., 2019).

18-day-old wild-type (WT) seedlings, grown under long-day photoperiod, 24°C, and 50% humidity conditions, were subjected to various abiotic stresses. The shoot outcomes of this assay were acquired from the Arabidopsis eFP Browser (http://bar.utoronto.ca/efp/cgi-bin/efpWeb.cgi) (Winter et al., 2007). Differential expression was determined by dividing the gene’s expression value under a specific abiotic stress condition by its corresponding control value, yielding the fold change of abiotic stress in comparison to the mock condition.

### In silico structural data analysis

Domains prediction was conducted using the InterPro tool (https://www.ebi.ac.uk/interpro/) (Merida et al., 2017)(Paysan-Lafosse et al., 2023). AlphaFold (https://alphafold.ebi.ac.uk/) (Jumper et al., 2021; Varadi et al., 2022) was used to predict the tertiary structure of DGK1, DGK2 and HsDGKε. The Intrinsically Disordered Region (IDR) domain of DGK1 and DGK2 was predicted using the program IUPRED3 (https://iupred3.elte.hu/plot).

### Transient expression in *Nicotiana benthamiana*

Different constructs were transformed into *Agrobacterium tumefaciens* (GV3101::pMP90) using electroporation, including p19 (to prevent gene silencing associated with overexpression). Subsequently, *A. tumefaciens* strains were utilized for the transient transformation of *N. benthamiana*. For this purpose, they were grown at 28°C overnight in Luria-Bertani (LB) medium supplemented with rifampicin (50 µg/ml), gentamicin (25 µg/ml), and the specific antibiotic corresponding to the construct (spectinomycin at 100 µg/ml or kanamycin at 50 µg/ml).

The cultures were centrifuged at 3,000 x g for 15 minutes at room temperature. The resulting pellets were resuspended in agroinfiltration solution [10 mM MES (pH 5.6), 10 mM MgCl2, and 1 mM acetosyringone], and incubated in the dark for 2 hours at room temperature. For single-gene expression experiments, the resuspended *Agrobacterium* cells were mixed to reach an OD_600_ of 0.70 for the construct and 0.25 for the p19 strain. In double infiltration experiments, *Agrobacterium* strains were used at OD_600_ of 0.40 for the constructs and 0.15 for the p19 strain. Two leaves from 3- week-old *N. benthamiana* plants (specifically, the 3rd and 4th leaves from the apex) were infiltrated on the abaxial side using a 1 ml syringe without a needle. The infiltrated plants were maintained under growth conditions for 2 days before the analysis by confocal microscopy and, if applicable, the sample collection.

### Confocal microscopy images

For confocal imaging, *N. benthamiana* leaves were infiltrated as described in “Transient expression in *Nicotiana benthamiana* ”. Leaf-disks were excised from the plants immediately before visualization. GFP, RFP or YFP fluorescence of the lower epidermis was visualized 2 days post infiltration by confocal microscopy. Confocal images were captured using the Zeiss LSM880 confocal microscope. The GFP and YFP excitation was achieved using the 488 nm argon laser, while for the RFP excitation the 561 nm laser was utilized. For co-localization, sequential line scanning mode was used to separate signals. Fluorophores detection involved a PMT and a GaAsp (used to improve the signal recognition), along with an additional PMT for transmitted light. Objectives employed were Plan-Apochromat 40X (water) and 63X (oil) with up to 4X digital zoom. Cortical plane images are a maximum Z projection of several planes (900 nm spacing) from the cell surface to the cell interior. The equatorial images used in the FRET assay correspond to single plane images. Microscopy image processing was performed using the program FIJI (Schindelin et al., 2012).

### BiFC Constructs for Expression in *N. benthamiana*

For Bimolecular fluorescence complementation (BiFC) assay, full-length cDNA of *DGK2* and *SYT1* were cloned into the pDEST-GW-VYNE and pDEST-GW-VYCE vectors (Gehl et al., 2009), respectively. Each vector contains one-half of YFP fused to the C-terminal of the target proteins (DGK2-nYFP and SYT1-cYFP). 35S:DGK2-nYFP and 35S:SYT1-cYFP expression vectors were transformed into *Agrobacterium* and infiltrated in *N. benthamiana* as detailed in the "Transient expression in *Nicotiana benthamiana*" section. Subsequently, the interaction between DGK2 and SYT1 was observed detecting, by confocal microscopy, the fluorescence produced by the binding of nYFP and cYFP, as described in “Confocal microscopy images”.

## FRET

Three-week-old *N. benthamiana* leaves, transiently co-expressing proteins tagged at the N-terminal to GFP and RFP, were used for Förster Resonance Energy Transfer (FRET) analyses. Confocal images were single plane captured using the Zeiss LSM880 confocal microscope with the Plan-Apochromat 40x/1.2 NA (water) objective lens. Equatorial sections of lower epidermal cells where both proteins co-localized with similar intensity values were examined. Three regions of interest (ROI) were measured: ROI 1 was photobleached over the acceptor fluorophore (RFP) using the 561 nm laser for 100 iterations at 100% power; ROI 2 was a random selected area not photobleached; and ROI 3 was background with no signal. In each ROI, six measurements of the donor fluorophore (GFP) were taken before photobleaching (Pre), and six measurements were taken after photobleaching (Post). The FRET efficiency was calculated as the percentage increase in donor fluorophore intensity (% ΔGFP) after acceptor removal using the formula: % ΔGFP = 100 x (Post-Pre)/Post (Liao et al., 2019). ROI 2 and 3 were technical control, where no increase of GFP intensity was found. ROI 1 data were used to calculate the interaction. More than 25 measurements in different cells, leaves and plants were taken for each protein pair. Similar results were obtained in three independent experiments.

### DGK1 segmentation and ER-PM CS quantification

Confocal images of the cortical plane of DGK1 transiently co-expressed in *N. benthamiana* leaves were segmented in ER and ER-PM CS in a semiautomatic way, using the interactive machine learning tool ilastik (Sommer et al., 2011). ER and ER-PM CS areas were quantified in FIJI (Schindelin et al., 2012) and the ratio ER-PM CS / ER was calculated for each ROI.

### Protein extraction and Immunoblot analysis

Arabidopsis or Nicotiana tissue was ground to a fine powder using liquid nitrogen. Proteins were extracted by incubating 100 mg of sample with Laemmli extraction buffer 2X [125 mM Tris-HCl, pH 6.8, 4% (w/v) SDS, 20% (v/v) glycerol, 2% (v/v) β-mercaptoethanol, and 0.01% (w/v) bromophenol blue] at 75°C for 30 minutes. Samples were then centrifuged for 1 minute at 20,000 g at room temperature, and the supernatants were collected.

Proteins were separated by size through polyacrylamide gel electrophoresis (SDS-PAGE). Subsequently, they were electrotransferred onto a polyvinylidene fluoride (PVDF) membrane (Immobilon-P, Millipore, 0.45 μm pore size, IPVH00010) following the manufacturer’s instructions. The membrane was blocked with 5% milk in Tris-buffered saline with Tween 20 (TTBS) for 2 hours at room temperature and then incubated at 4°C overnight with TTBS 1% milk and the appropriate primary antibody [anti-GFP 1:600 (Santa Cruz Biotechnology, sc-9996); anti-RFP 1:2,000 (ChromoTek, 6g6); anti-AHA3 1:10,000 (a gift from Ramón Serrano); anti-BIP 1:2,500 (Agrisera, AS09 481); Sec 21P 1:1,1000 (Agrisera, AS08 327); V-ATPasa 1:2,000 (Agrisera, AS07 213); anti-TOC75-3 1:2,000 (Agrisera, AS08 351)]. Later, the membrane was incubated for 2 hours at room temperature with TTBS 1% milk and the corresponding secondary antibody conjugated to horseradish peroxidase [anti-mouse 1:80.000 (Sigma-Aldrich, A9044) or anti-rabbit 1:80,000 (Sigma-Aldrich, A0545)]. Proteins and epitope-tagged proteins were detected in the membrane using the ChemiDoc XRS+ imaging system (BioRad) with either Clarity Western ECL reagent (170-5060) from BioRad or SuperSignal West Atto (A38555) from Thermo Scientific for enhanced sensitivity detection. Only images with no saturated pixels were used for protein quantification in FIJI (Schindelin et al., 2012). Following immunodetection, the membrane was stained with Coomassie (R-250) to confirm uniform loading.

### Yeast Two-Hybrid Screening

Yeast two-hybrid screening was conducted by Hybrigenics Services, S.A.S., Paris, France (http://www.hybrigenics-services.com). The C2 domain of SYT1 coding region (Leu244 to Ser541) was PCR amplified and inserted into pB27 as a N-terminal fusion to LexA DNA-binding domain (SYT1C2-LexA). The construct was confirmed by sequencing and used as a bait to screen a random-primed cDNA library of 1 week-old seedlings Arabidopsis. After library screening, 17 colonies were selected on a medium lacking tryptophan, leucine and histidine. The prey fragments of the positive clones were amplified by PCR and sequenced at their 5′ and 3′ end. Of these colonies, 12 independent clones were identified in the screening with only one had a good Predicted Biological Score. This clone corresponded to the amino acids 231-1546 of the DGK2 (AT5G63770).

### Co-IP assay

Co-Immunoprecipitation (Co-IP) experiments were conducted following the procedures outlined in Amorim-Silva et al. (2019) with slight adjustments. 3-week-old *N. benthamiana* leaves, transiently transformed, were ground. Proteins were extracted from 0.5 g of tissue per sample by incubating the powder for 30 minutes on an end-over-end rocker at 4°C with 1 ml of non-denaturing extraction buffer [50 mM Tris–HCl, pH 7.5, 150 mM NaCl, 1% (v/v) Nonidet P-40, 10 mM EDTA, 1 mM Na2MoO4, 1 mM NaF, 10 mM DTT, 0.5 mM PMSF, and 1% (v/v) protease inhibitor (Sigma, P9599)]. The extract was then centrifuged at 15,000 g for 20 minutes at 4°C and filtered by gravity through Poly-Prep chromatography columns (Bio-Rad, 731-1550). At this stage, 100 μL of the supernatant was saved as input for Western Blot, and the remaining portion was incubated for 2 hours with 30 μL of GFP-fused protein agarose beads (Chromotek). During incubation of protein samples with beads, the final concentration of detergent (Nonidet P-40) was adjusted to 0.2% (v/v) in all cases to avoid unspecific binding to the matrix as recommended by the manufacturer. Beads were washed four times with the extraction buffer without detergent, resuspended in 75 μL of 2X Laemllie Buffer and heated at 75°C for 30min to dissociate immunocomplexes from the beads. Once the proteins were purified, the samples were further processed as described in the "Protein extraction and Immunoblot analysis" section, with both Input and immunoprecipitated (IP) samples run in duplicate SDS-PAGE gels for detection with both GFP and RFP antibodies. Co-IP assays were performed twice with equal result.

### Phylogenetic analysis

The protein sequences of all human and Arabidopsis DGKs were obtained from the UniProt database (https://www.uniprot.org/), the alignment was done by Clustal W (https://www.ebi.ac.uk/Tools/msa/clustalo/) and phylogenetic analyses were conducted using the software MEGA X (Kumar et al., 2018). The evolutionary history was inferred by using the Maximum Likelihood method and JTT matrix-based model. The bootstrap consensus tree inferred from 500 replicates is taken to represent the evolutionary history of the taxa analyzed. Branches corresponding to partitions reproduced in less than 50% bootstrap replicates are collapsed. The percentage of replicate trees in which the associated taxa clustered together in the bootstrap test (500 replicates) are shown next to the branches. Initial tree for the heuristic search was obtained automatically by applying Neighbor-Join and BioNJ algorithms to a matrix of pairwise distances estimated using a JTT model, and then selecting the topology with superior log likelihood value. This analysis involved 17 amino acid sequences and there was a total of 344 positions in the final dataset.

### Analysis of disorder and prion-like regions

Prion-like residues of DGK1 and DGK2 were obtained using the PLAAC online tool with the defective values, i.e. core length of 60 and a relative weighting of 100. Proteins regions were considered disordered if they had a MobiDB consensus score above 0.15 (15% disordered residues), and prion-like if they contained a prion-domain (≥ 60 amino acids).

#### RNA extraction for RT-PCR and RNAseq

Total RNA extraction was conducted using the E.Z.N.A Plant RNA kit (BIO-TEK, R6827-01) in accordance with the manufacturer’s instructions. For this purpose, 50 mg of tissue ground with liquid nitrogen was used. DNase treatment was performed on the extracted RNA to eliminate any genomic remnants using the RNase-free DNase I Set (Omega BIO-TEK, E1091). RNA purity, integrity, and concentration were assessed using Nanodrop One and by running samples on agarose gel.

For RT-PCR analysis, complete 10-day-old seedlings grown on plates were used as the tissue source. 500 μg of RNA were reverse transcribed into complementary DNA (cDNA) using the iScript reverse transcriptase (1708890, BioRad) following instructions by the manufacturer. The absence of undesired genomic residues in the RNA and the successful conversion to cDNA were verified by LOX2 diagnostic PCR. LOX2 was not amplified in the RNA samples but was amplified in the cDNA samples. RT-PCR was performed with specific primers (Supplemental Table S5) and it was observed that in both *dgk1-1* and *dgk2-2* mutants, the T-DNA interruption led to transcriptional disruption.

For RNA sequencing (RNAseq), the aerial parts of a pool of 2-week-old seedlings (10 seedlings per biological replicate, and at least 3 replicates per group) grown in soil under control condition were collected.

### RNAseq bioinformatic analysis

The clustering of the index-coded samples was performed on a cBot Cluster Generation System using TruSeq PE Cluster Kit v3-cBot-HS (Illumina) according to the manufacturer’s instructions. After cluster generation, the library preparations were sequenced on an Illumina Novaseq platform and 150 bp paired-end reads were generated at the Novogenelll(UK) Company Ltd. At last, RNA integrity was assessed using the RNA Nano 6000 Assay Kit of the Bioanalyzer 2100 system (Agilent Technologies, CA, USA). The reads were quality filtered and trimmed using Trimmomatic version 0.36 (Bolger, Lohse, & Usadel, 2014) with default paired-end mode options. The resulting reads were then aligned to the TAIR10 version of the Arabidopsis thaliana genome sequence (https://www.arabidopsis.org/) using Hisat2 version 2.1.0 (Kim, Langmead, & Salzberg, 2015). These read alignments (in BAM format) were used for transcript quantification with the cuffdiff program of the Cufflinks version 2.2.1 package (Trapnell et al., 2013). The resulting read alignments were visualized and clustered using Tablet software (Milne et al., 2013) and CummeRbund R package version 2.23.0 (Goff, Trapnell, & Kelley, 2014). To determine DEGs, a False Discovery Rate (q-value) cutoff of ≤ 0,05 was set. DEGs were subjected to singular enrichment analysis for the identification of overrepresented GO terms using PANTHER (Thomas et al., 2022) with the default options (Fisher’s Exact as the test type and the Bonferroni correction for multiple testing, p < 0.05). The RNA seq data from this manuscript have been submitted to Gene Expression Omnibus (GEO; https://www.ncbi.nlm.nih.gov/geo/) and assigned the identifier PRJNA1062618.

### Metabolomic Analysis

The samples used for the transcriptomic analysis were also used for metabolomic analysis (2-week-old Arabidopsis seedlings grown in soil under standard conditions). 50 mg of powder were analyzed by the metabolomics service at IHSM following the adapted version of the method described in Osorio et al., 2012. The equipment used was Agilent 7890B gas chromatograph system coupled with a Pegasus® HT High Throughput TOF-MS spectrometer (GC-TOF/MS; Agilent Technologies Inc. CA, USA).

### Plasma membrane and Inner Membranes isolation

For membrane isolation, WT and *dgk1dgk2* Arabidopsis plants were initially grown vertically on plates under control conditions with a short-day photoperiod for 7 days. Subsequently, the seedlings were transferred to soil and kept under the same conditions for an additional week before transitioning to a long-day photoperiod for 3 weeks to encourage larger rosette growth. Just before bolting, the plants were subjected to a 3-day cold treatment at 4°C while maintaining the photoperiod. The aerial parts were then harvested for the Plasma Membrane and Inner Membranes isolation, that was carried out as described previously in Bernfur et al., 2013 with some modifications. In brief, five grams of 5-week-old Arabidopsis leaves were homogenized using an Ultraturrax in 35 ml of homogenization buffer [330 mM sucrose, 50 mM MOPS-KOH, pH 7.5, 5 mM EDTA, 5 mM EGTA, 20 mM NaF, 5 mM ascorbate, 5 mM DTT, 150 μM Protease Inhibitor (Pefabloc, 11429868001), and 0.6% (w/v) polyvinylpolypyrrolidone (PVPP)]. DTT and Protease Inhibitor were added just before use. All steps of this protocol were performed at 4°C to prevent degradation. The homogenate was filtered through 4 layers of nylon mesh (200 μm) and centrifuged at 10,000 g for 15 minutes to precipitate and discard non-homogenized cellular debris. The remaining fraction (Total, T) was centrifuged at 100,000 g in a swinging JS24.38 rotor for 2 hours at 4°C. The supernatant was discarded, and the precipitated microsomal fraction (M) was resuspended in resuspension buffer (0.33 M sucrose, 5 mM K-phosphate at pH 7.8, 0.1 mM EDTA, and 1 mM DTT) to a final weight of 6 g. The aqueous polymer two-phase partitioning protocol consists of 4 systems, each containing: 6.1% (w/w) Dextran T500, 6.1% (w/w) PEG 3350, 330 mM sucrose, 5 mM K-phosphate pH 7.8, 3 mM KCl, and distilled water up to 9 g. To two of the 4 systems, 3 g of the microsomal fraction were added to form a 12 g system, and to the other 2 systems, 3 g of resuspension buffer were added. Following the protocol by Larsson et al., 1994 2 phases enriched in Plasma Membranes (PM) and 2 in Inner Membranes (IM) were obtained, combined, and diluted 3 times in resuspension buffer before being centrifuged at 100,000 g for 2 hours at 4°C. The PM pellet was resuspended in 100 μL of resuspension buffer supplemented with 5 mM KCl, and the IM pellet was resuspended in 1 ml of the same buffer. Samples were stored at -80°C until lipid extraction.

### Lipid extraction and analysis

Lipids were extracted from 100 μL of PM or IM. Glassware was used throughout the procedure. The samples, along with 1 ml of isopropanol, were incubated at 75°C for 20 minutes. Subsequently, 2 ml of chloroform/methanol (1:1) and 0.7 ml of water were added. After 30 seconds of vortex, an additional 2 ml of chloroform/water (1:1) was added. The mixture was centrifuged for 3 minutes at 500 g, and the lower chloroform phase was gently transferred to a new tube. A re-extraction of lipids was performed by adding 1 ml of chloroform to the first tube, repeating the centrifugation, and recovering the lower phase. Both extractions were combined. The chloroform was evaporated using N_2_ gas while keeping the sample in a 37°C block. Once all the solvent was evaporated, the lipids were resuspended in 200 μL of chloroform and stored at -80°C for subsequent analysis.

Analysis of lipids including neutral (DAGs) and polar lipids (PC, PE, PI, PG, lysophosphatidylcholine [LPC], MGDG, and DGDG) were carried out using high-resolution/accurate mass (HR/AM) lipidomics with a Vanquish - Q Exactive Plus UPLC-MS/MS system (Thermo Fisher Scientific). The analytical protocol is based on the publication by Bird, SS., Marur,VR, Sniatynski, MJ., Greenberg,HK., and Kristal, BS. Lipidomics Profiling by High-Resolution LC−MS and High-Energy Collisional Dissociation Fragmentation: Focus on Characterization of Mitochondrial Cardiolipins and Monolysocardiolipins, Analytical Chemistry 83 (3), 940-949 (2011) https://doi.org/10.1021/ac102598u with some modifications. Briefly 10µl total lipid extract was injected into the UPLC/MS (Thermo Vanquish system). Separation occurred on a Thermo Scientific Accucore C18 column (2.1 x 150 mm, 2.6 mm) at 35°C, with the autosampler tray temperature set at 10°C and a flow rate of 400μl min^-1^. The mobile phase consisted of A: 10 mM ammonium formate in 50% acetonitrile plus 0.1% formic acid and B: 2 mM ammonium formate in acetonitrile/propan-2-ol/water 10/88/2 plus 0.02% formic acid. The elution gradient spanned 28 minutes, starting at 35% B and reaching 100% at 24 minutes.

The Thermo Q Exactive HESI II conditions utilized a sweep plate in the C probe position. Conditions were adjusted for separate positive and negative runs, with running samples in a single polarity resulting in more identifications. LC/MS was performed at 140K resolution, and HCD MS2 experiments (35K resolution) were conducted in positive and negative ion modes. Full scans were conducted at 140,000 resolution from m/z 150-1200, followed by top 15 MS/MS at 35,000 resolution. The stepped collision energy was set at 25, 30, and 40, with the replacement of 25 with 30 in negative ion mode.

For positive ion mode, sheath gas was set to 60, Aux gas to 20, sweep gas to 1, and spray voltage to 3.2 KV with slight adjustments in negative ion mode. Capillary temperature was maintained at 320oC, and the aux gas heater was set to 370oC. Lipid search 5 by Thermo Fisher Scientific was used for lipid characterization. Potential lipid species were identified separately from positive or negative ion adducts. The data from each replicate were aligned within a chromatographic time window by combining positive and negative ion annotations, subsequently merged into a single lipid annotation. Identified lipid molecular species were quantified using polar and neutral lipid standards, 13:0-LPC, di24:1-PC, di14:0-PE, di18:0-PI, di14:0-PG, di18:0-PS, and di14:0-PA (supplied by Avanti Polar Lipids, USA), 0.857 nmol of tri15:0 TAG and 0.043 nmol 18:0-20:4 DAG (supplied by Nu-Chek-Prep). Full documentation of lipid profiling data is provided in Supplemental Table S4.

### Cold assays for measuring root elongation

Freshly harvested seeds of different genotypes, including wild type (WT), mutants, and complementation lines, were plated as described in “Plant manipulation and growth conditions”. Seed were vertically grown on plates with half-strength MS and 1.5% sucrose under control temperature (22°C ± 1°C) and light conditions (long-day photoperiod). Equivalent three-day-old seedlings were transferred to new plates that were randomly divided between control and cold. The control corresponds to plants kept for two weeks at 22°C ± 1°C, and the cold treated plants were moved to growth chambers at 10°C with the same photoperiod for four weeks. After this period, seedlings were photographed and root length were measured using FIJI software (Schindelin et al., 2012). Three independent experiments showed similar results.

### Freezing assays

Freezing tolerance assays were conducted on soil-grown plants in growth chambers at 20°C under long-day conditions for two weeks. To acclimate them to cold temperatures, the plants were transferred to growth chambers at 4°C under long-day photoperiod with a light intensity of 40 μmol m^-2^ s^-1^ for 7 days. Freezing treatments were performed by keeping the plants at 4°C for one hour, after which the temperature was lowered (1 °C / 30 minutes) until a temperature of -10°C was reached. After 6 hours at -10°C, the temperature was elevated to 4°C (1°C / 30 minutes). Subsequently, the plants were returned to growth chambers at 20°C under long-day conditions, and after one week of recovery, survival rates were assessed. Five biological replicates (> 20 plants per replicate) were performed.

### Statistic and graphs

RNAseq overlap enrichment between *dgk1dgk2* and *syt1* and the associated p-value were calculated by using the Web http://nemates.org/MA/progs/representation.stats.html. Enrichment was calculated as the number of overlapping genes divided by the expected number of overlapping genes drawn from both independent groups. The expected genes were defined by next formula: (n° of genes in *dgk1dgk2* * n° of genes in *syt1)* / total genes analyzed in full genome.

Statistical analyses were conducted using Prism 8.02 for Microsoft (GraphPad Software, www.graphpad.com). The tests performed included unpaired t-tests (*p < 0.05), one-way or two-ways analysis of variance (ANOVA) followed by Tukey’s multiple comparison test or Dunnett’s multiple comparisons test (****p < 0,0001; ***p < 0,0002; **p < 0,0021; *p < 0,0332), respectively. In the figures, asterisks denote statistical variations between the mutant and Col-0, or between the treatment and the mock. Different lowercase letters in the graphs indicate significant differences. The data represents the mean values, with error bars depicting standard deviation (SD). In the figure legends, “n” represents the number of plants phenotypically analyzed or the number of ROI analyzed for FRET quantification or ER-PM CS/ER ratio in segmentation. The experiments were repeated a minimum of three times, yielding similar results. Graphs were performed mainly in Prism 8.02 and bubble plots in Prism 9.00.

### Accession numbers

The genes investigated in this research are catalogued in The Arabidopsis Information Resource (https://www.arabidopsis.org/) with the corresponding accession numbers: *DGK1*: AT5G07920; *DGK2*: AT5G63770; *DGK3*: AT2G18730; *DGK4*: AT5G57690; *DGK5*: AT2G20900; *DGK6*: AT4G28130; *DGK7*: AT4G30340; *SYT1*: AT2G20990.

## Acknowledgements

We would like to thank Jorge Morello López, for technical assistance in the use of the Ilastik tool, Alicia Esteban del Valle, for confocal images assistance, and Tabata Rosas for providing the FRET protocol, all from IHSM-CSIC-UMA (Instituto de Hortofruticultura Subtropical y Mediterránea, Málaga, Spain). We also thank Susana Silvestre for helping in the management of sending samples. We appreciate the equipment provided by Remedios Crespillo and the SCAI Facility at the University of Málaga. The ER marker was a gift from Iraida Amaya Saavedra (IFAPA-Centro de Churriana, Málaga, Spain). We thank Plan Propio from the University of Málaga for financial support.

---

**Figure S1.**
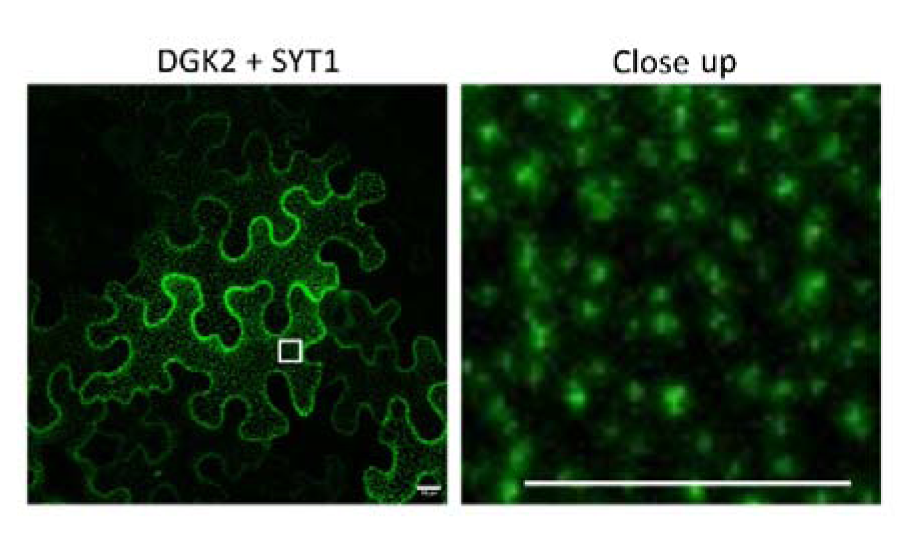
SYT1 interacts with DGK2 at ER-PM CS. The punctate pattern observed in the BiFC assay when SYT1 and DGK2 are co-expressed in *N. benthamiana* indicates that its interaction occurs in ER-PM CS. Images were taken 2 days post-inoculation from the cortical region of lower epidermis cells. Scale bar = 10 µm.

**Figure S2.**
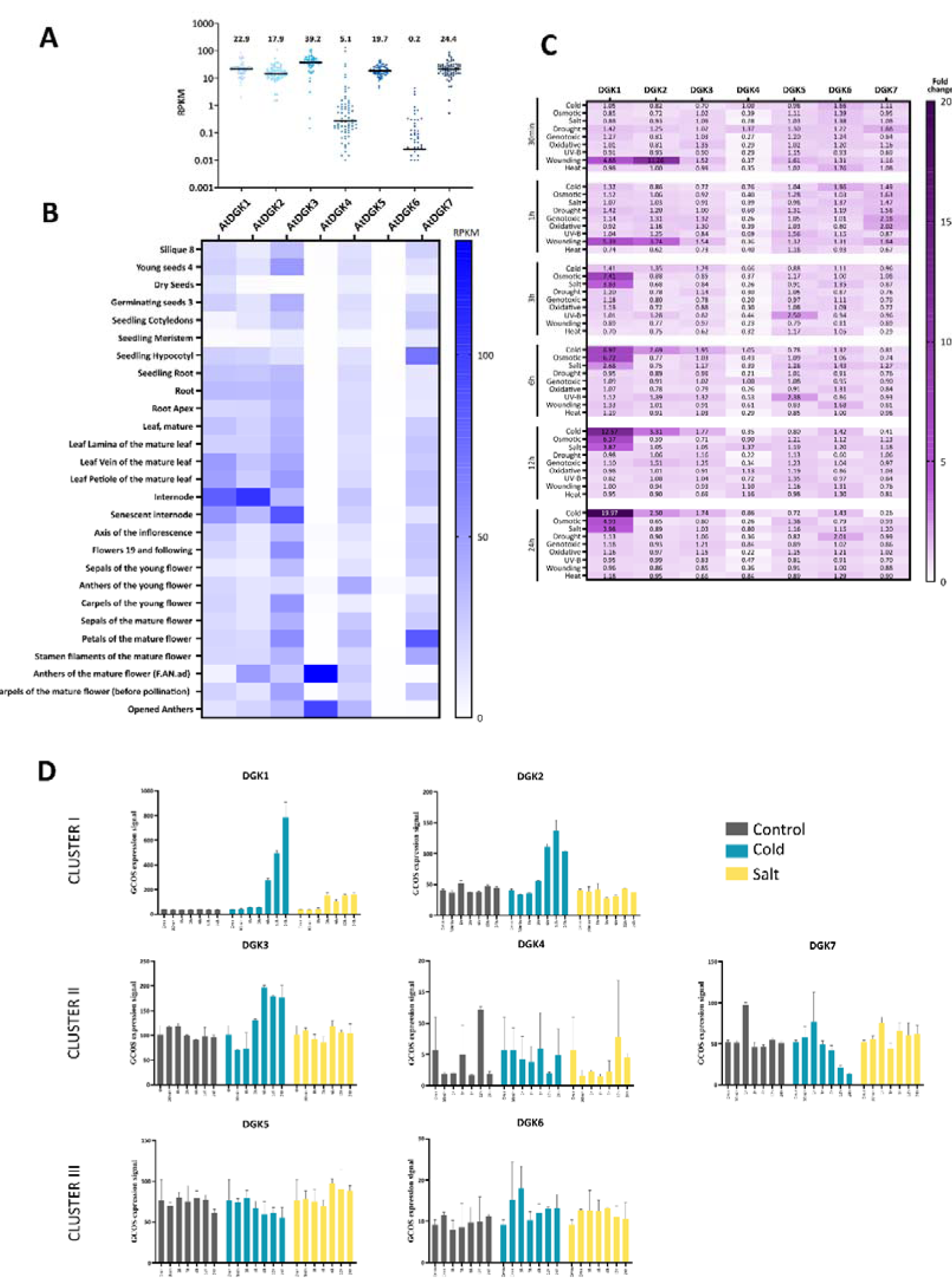
*DGKs* expression in different tissues and under several stresses. *DGK1* and *DGK2* are the most activated under abiotic stress. **A and B)** RNAseq data of *DGK1, DGK2, DGK3, DGK4, DGK5, DGK6* and *DGK7* obtained from vegetative tissues at different developmental stages from eFP-seq Browser (https://bar.utoronto.ca/eFP-Seq_Browser/). A) Schematic summary of the level of expression of *DGK* genes. Each dot represents a value of RPKM and the bar represent the median, whose value is written above. B) Tissue expression pattern is represented by Heatmap, where white indicates the lowest expression (0 RPKM) and dark blue the highest value (132 RPKM), which corresponds to *DGK4* in anthers. **C and D)** Transcriptional response of *DGK* to abiotic stress. Data were collected from the eFP browser (http://bar.utoronto.ca/efp/cgi-bin/efpWeb.cgi?dataSource=Abiotic_Stress). C) Heatmap representing the fold change of DGK expression levels at 30 min, 1 h, 3 h, 6 h, 12 h, and 24 h after treatment compared to control (time 0 min, before treatment). D) *DGK* expression patterns under cold (4°C) and salt (150 mM NaCl) stress show *DGK1* and *DGK2* as the main *DGKs* activated under these abiotic stress conditions, mainly in cold.

**Figure S3.**
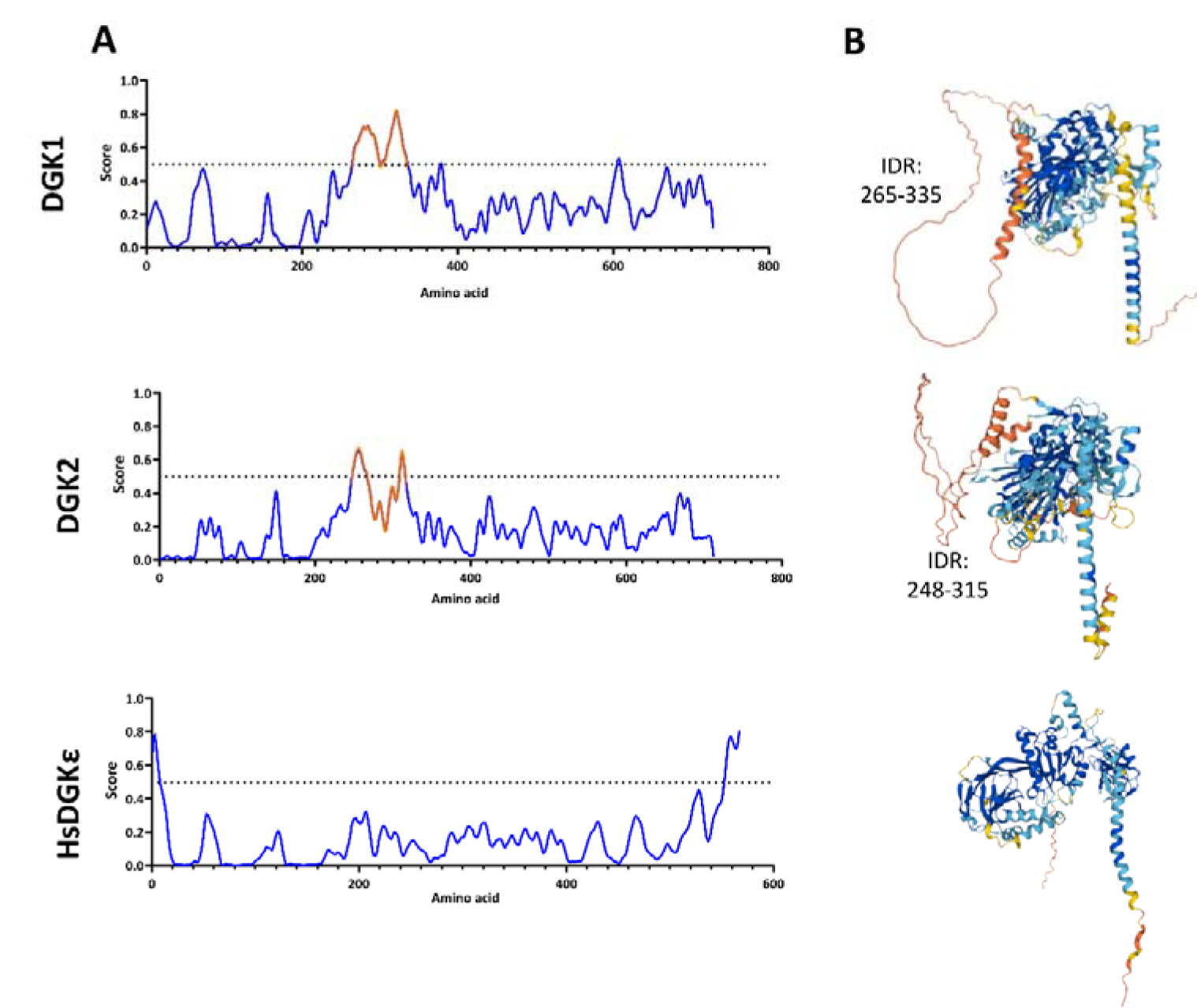
DGK1 and DGK2 feature an IDR between the C1 and Catalytic domains, whereas HsDGKε does not. A) The tool used to identify regions of disorder (https://iupred3.elte.hu/plot) showed the presence of IDRs in DGK1, DGK2 and HsDGKε. The score indicates the probability that each residue is part of a disordered region. Regions with a higher probability of being IDR are indiacted in orange. **B)** AlphaFold’s prediction, represented by UniProt, displays a per-residue confidence score. In this representation, the lowest scores are shown in orange, while the highest scores appear in blue. DGK1 and DGK2 regions with the lowest confidence score correspond to the IDR identified in panel A.

**Figure S4.**
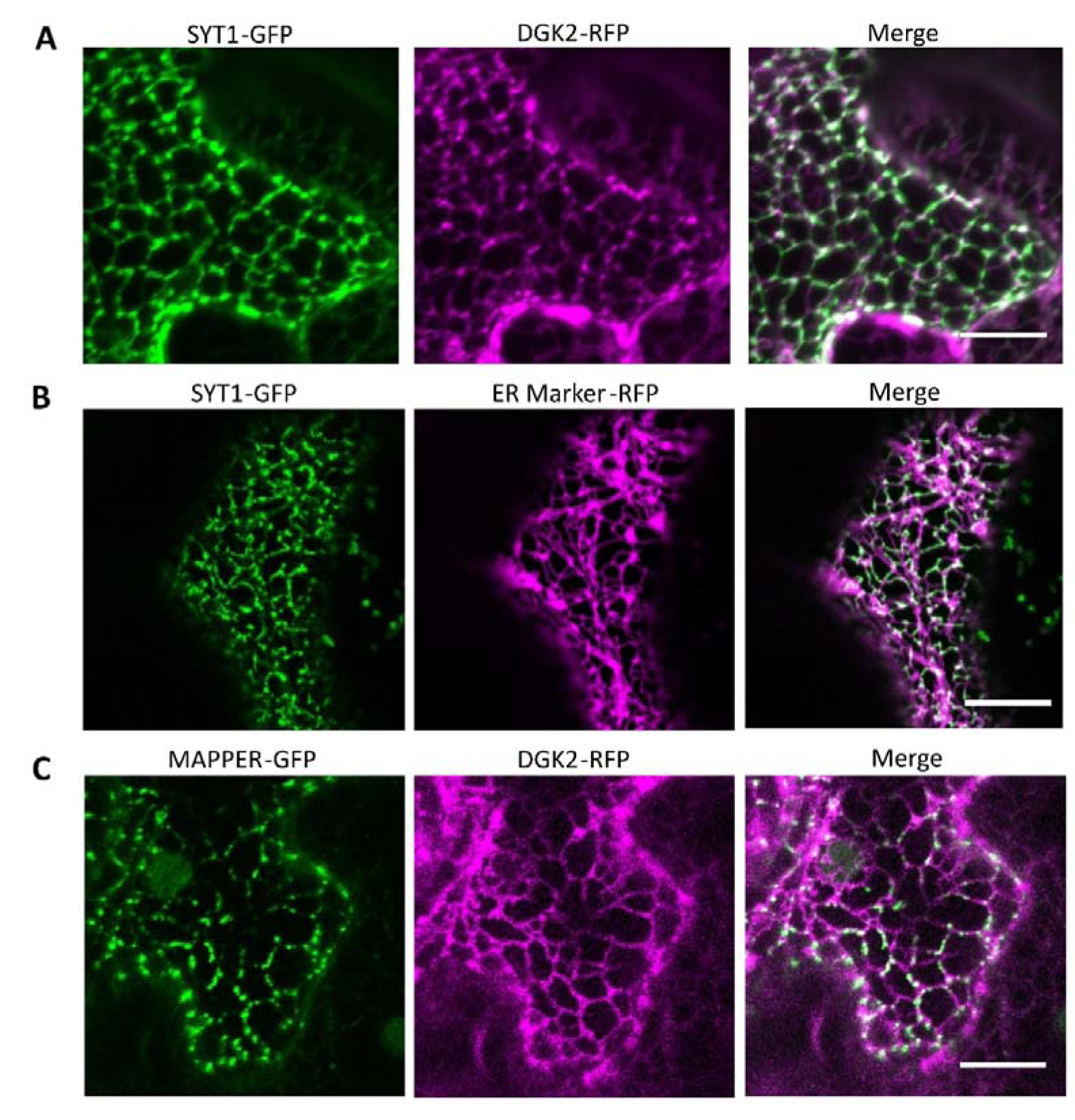
DGK2, as DGK1, is relocalized to ER-PM CS when SYT1 is co-expressed. A to C) *N. benthamiana* leaves co-expressing SYT1-GFP with DGK2-RFP (A) or with an ER Marker-RFP (B) show that SYT1 drags DGK2 but not the ER Marker (FaFAH) to the ER-PM CS. Additionally, the co-expression of DGK2-RFP with MAPPER-GFP does not change the reticular pattern of DGK2 (C). Confocal images were taken in the cortical region of lower epidermis cells 2 days post-infiltration. Scale bar = 10 µm.

**Figure S5.**
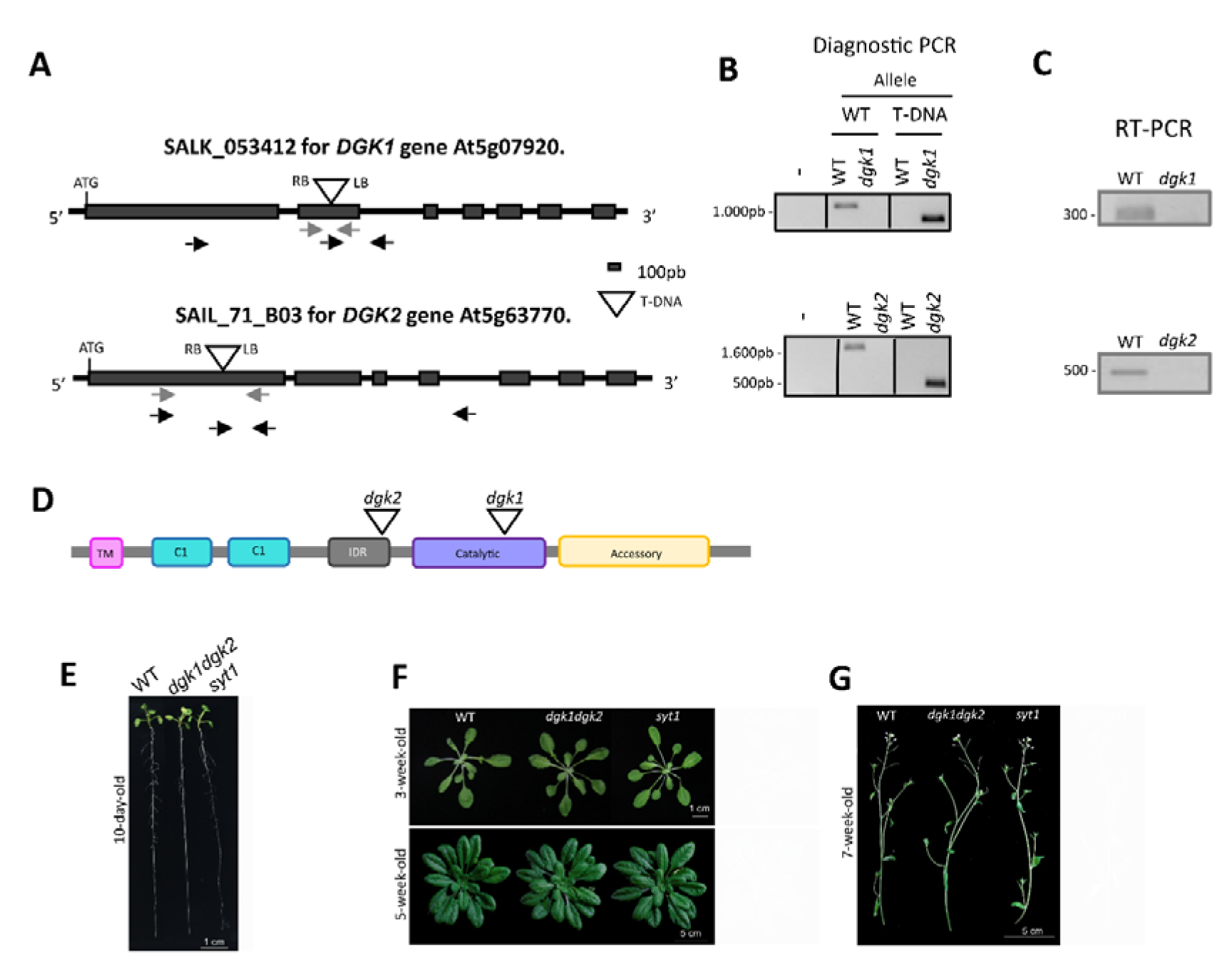
The mutants *dgk1dgk2* and *syt1* do not exhibit an apparent phenotype under control conditions. **A)** Schematic representation of the genomics of *DGK1* (At5g07920) and *DGK2* (At5g63770). The position of the start and stop codons are indicated. T-DNA insertions in SALK_053412 for *DGK1* and SAIL_71_B03 for *DGK2* are represented by triangles. LB, left border of inserted T-DNA; RB, right border of T-DNA; box, exon; line, intron; Black arrows, primers used for diagnostic PCR (B). Gray arrows, primers used for RT-PCR (C). **B)** Homozygous mutants for *dgk1-1* and *dgk2-2* were confirmed using diagnostic PCR with two pairs of oligos, one for the wild-type (WT) allele and the other for detecting the T-DNA insertion. **C)** The expression of *dgk1-1* (SALK_053412) and *dgk2-2* (SAIL_71_B03) was checked by RT-PCR to confirmed that both are Knock-Out mutants. **D)** Scheme of DGK1/2 protein domains. The T-DNA has been represented as a triangle in the corresponding insertion site in the genome. **E - G)** WT, *dgk1dgk2 and syt1* do not exhibit different phenotypes under any growth state when grown under control conditions. E) 10-day-old seedlings grown on vertical plates. F) Rosettes of 3 and 5 weeks grown in soil. G) Internodes of 7-week-old plants. Scale is indicated in the figure.

**Figure S6.**
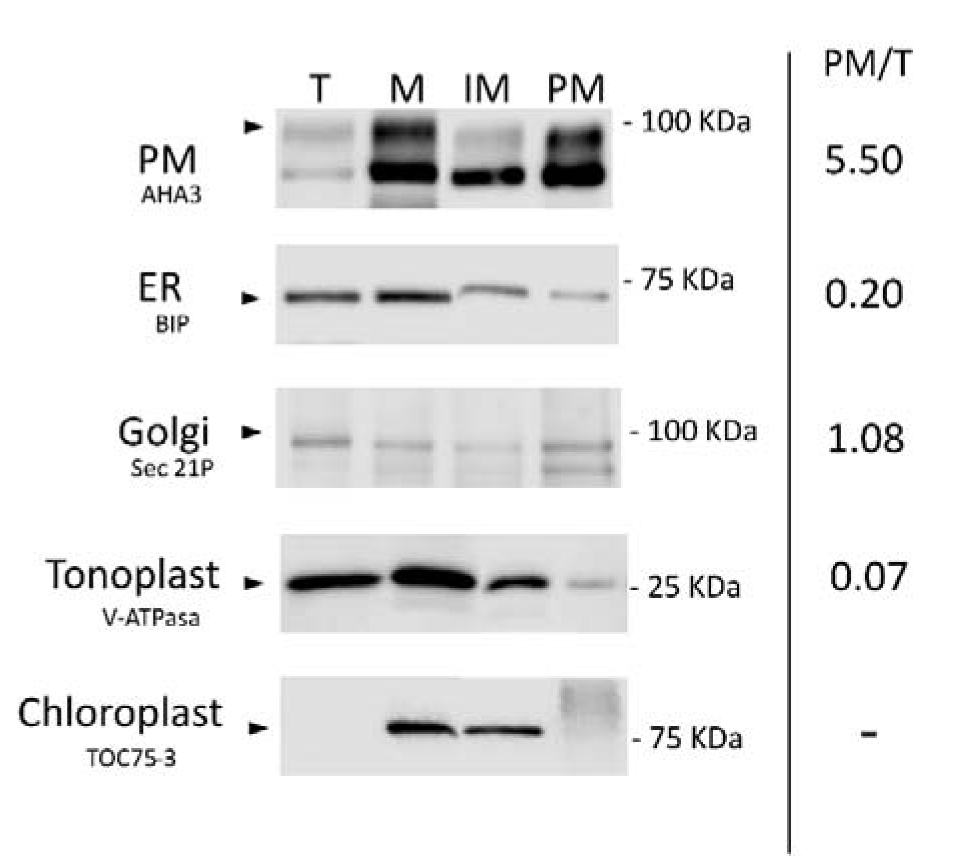
Inner Membranes and Plasma Membrane isolation and purity determination. Aerial part from 5-week-old Arabidopsis plants were used for the Plasma Membrane and Inner Membranes isolation. Samples from Total (T) fraction, which is the crude unpurified extract, microsomes (M), Inner Membrane (IM), and Plasma Membrane (PM) fractions were analyzed by immunoblotting using the organelle-specific antibodies indicated in ‘Methods’. Quantification of the different organelles shows that PM is highly enriched in PM fraction compare to the Total extract, while other membranes, as ER or tonoplast, decrease.

**Figure S7.**
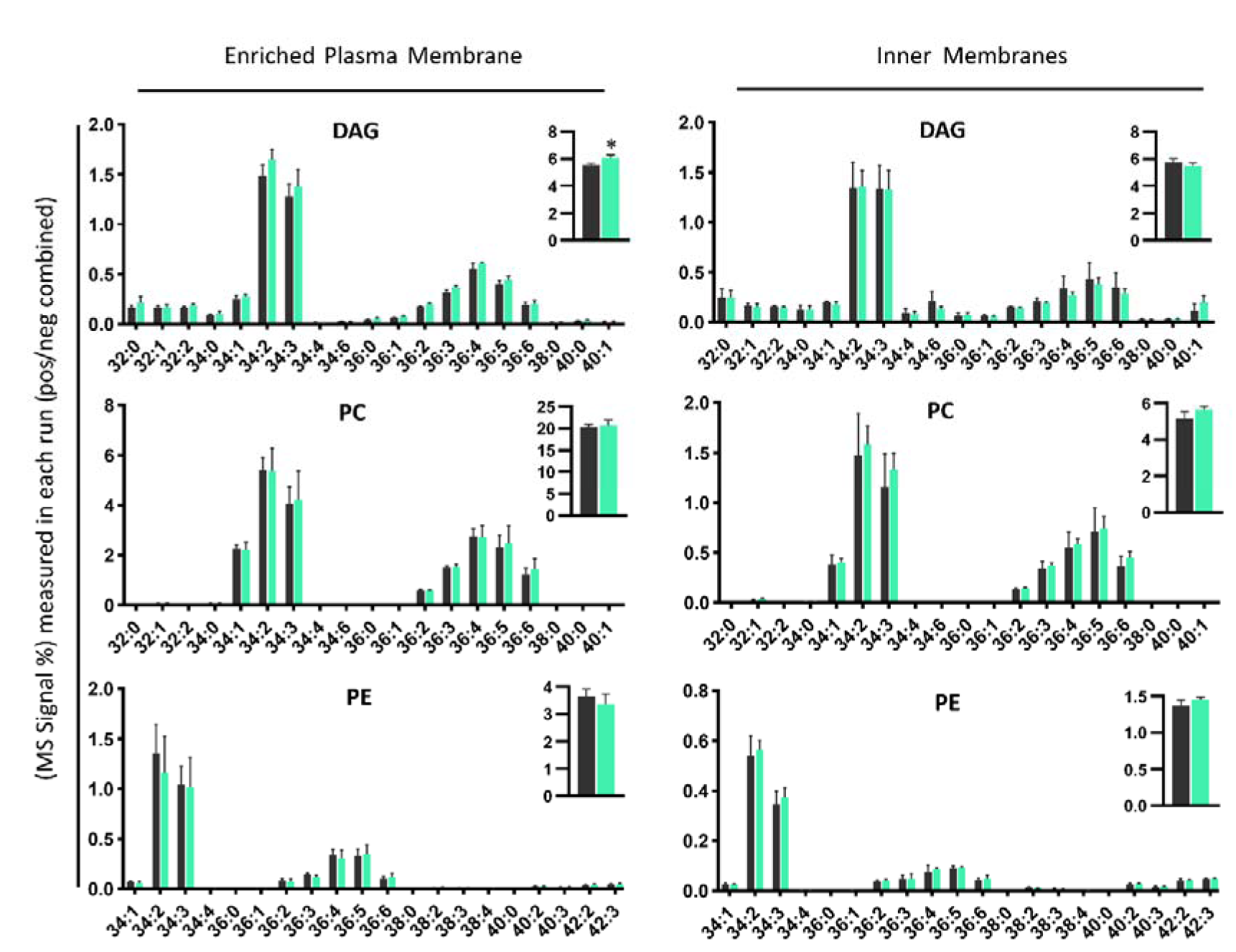
DAG accumulates slightly at the Inner Membrane (IM) in the *dgk1dgk2* mutant in Control conditions. High-resolution/accurate mass (HR/AM) lipid analysis of the molecular species of DAG, PC and PE in Plasma Membrane (PM) and Inner Membranes (IM). Fractions were isolated from 5-week-old WT and *dgk1dgk2* rosettes grown at control condition. PM and IM samples were purified by two phase partitioning protocol and lipids were extracted following as described in ‘Methods’. Acyl chains are expressed as number of acyl carbons: number of acyl double bonds. Distribution of the identified DAG, PC, and PE molecular species in the PM and in the IM of WT (gray) and *dgk1dgk2* (green) is represented. Column bars show the mean values of at least three biological replicates, with error bars indicating the standard error of the mean (SEM). The asterisks indicate statistically significant differences between *dgk1dgk2* and WT as determined by a Dunnett’s multiple comparisons test: ****p < 0,0001; ***p < 0,0002; **p < 0,0021; *p < 0,0332.

**Figure S8.**
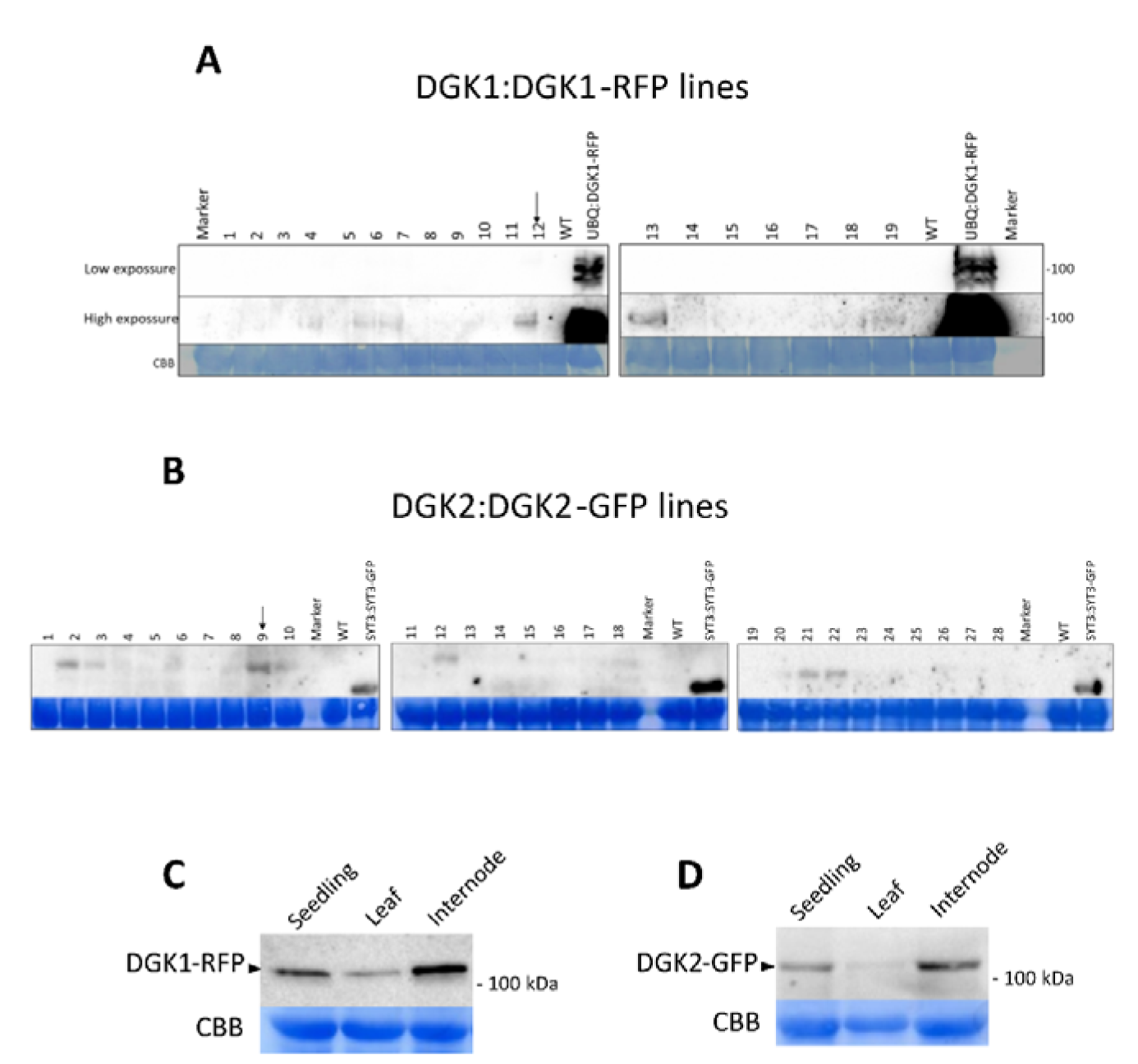
Immunoblotting assay of DGK1 and DGK2 Arabidopsis transgenic lines expressed under their endogenous promoters. Anti-RFP and anti-GFP antibodies were employed. **A and B)** More than 20 lines of DGK1:DGK1-RFP (A) and DGK2:DGK2-GFP (B) were generated and analyzed by Western Blot as described in ‘Protein extraction and Immunoblot analysis‘. The tissue used was a pool of T2 2-week-old seedlings grown in plates. The control UBQ:DGK1-RFP, expressed transiently in *N. benthamiana*, is detected with low exposure, while the Arabidospsis transgenic lines driven by their own promoter need high expossure to be detectable. Arrows indicate the lines that were later used in expression assays. **C and D)** Tissue-specific pattern of DGK1 (C) and DGK2 (D) protein amount in 2-week-old seedling, adult leaf and internode (7 weeks) grown in control conditions.

**Figure S9.**
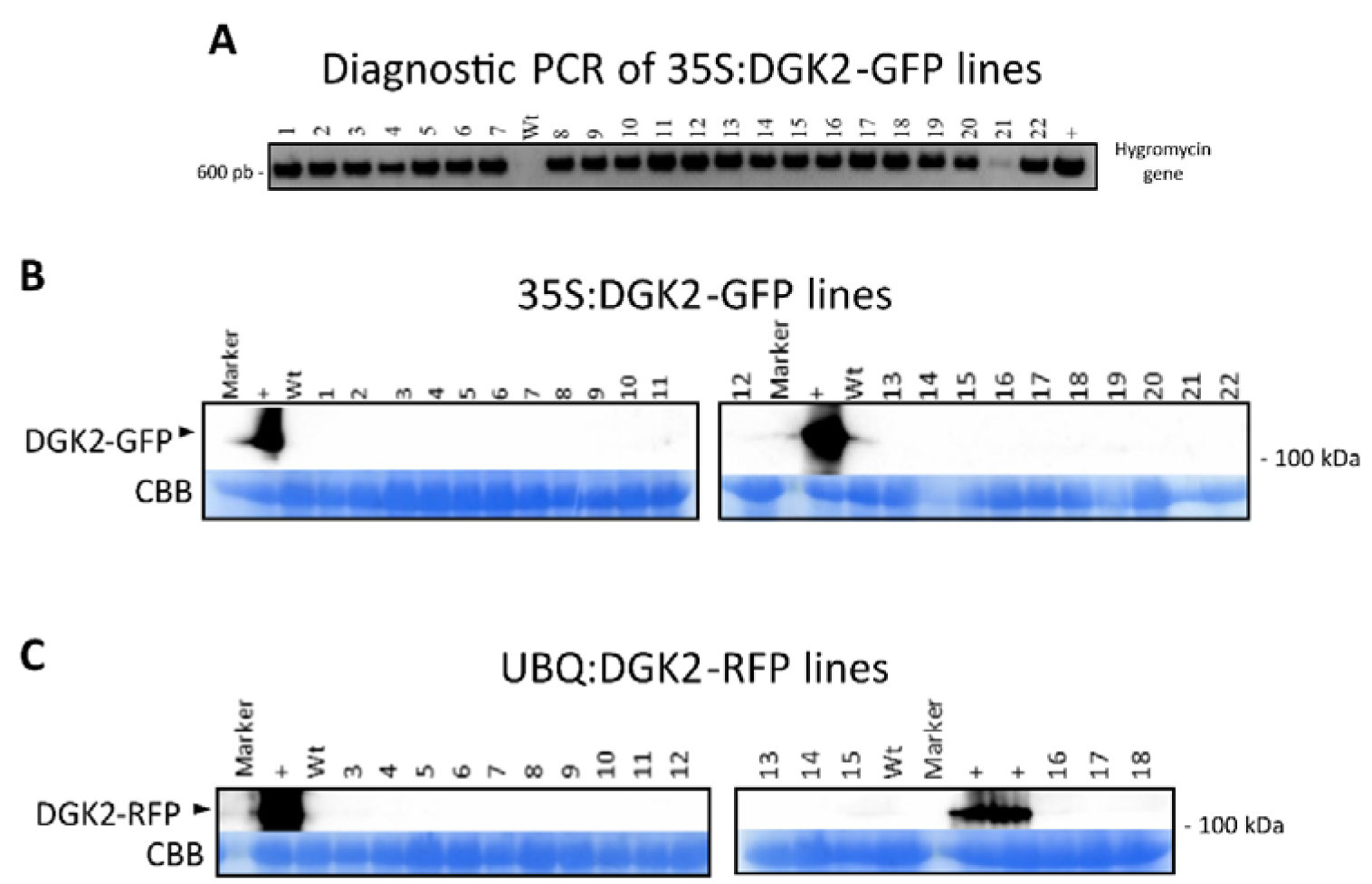
Overexpression of DGK2 causes its silencing. **A)** Diagnostic PCR of the hygromycin resistance gene, which was transformed together with the 35:DGK2-GFP construct, in all analyzed lines. **B and C)** Immunoblot assays in 35S:DGK2-GFP (B) and UBQ:DGK2-RFP (C) Arabidopsis lines. There was no detectable signal in any of them. Positives samples were the same constructs expressed transiently in *N. benthamiana* . A pool of T2 2-week-old seedlings grown in plates were used to determine the DGK2 expression by Western Blot. Anti-RFP and anti-GFP antibodies were employed.

**Figure S10.**
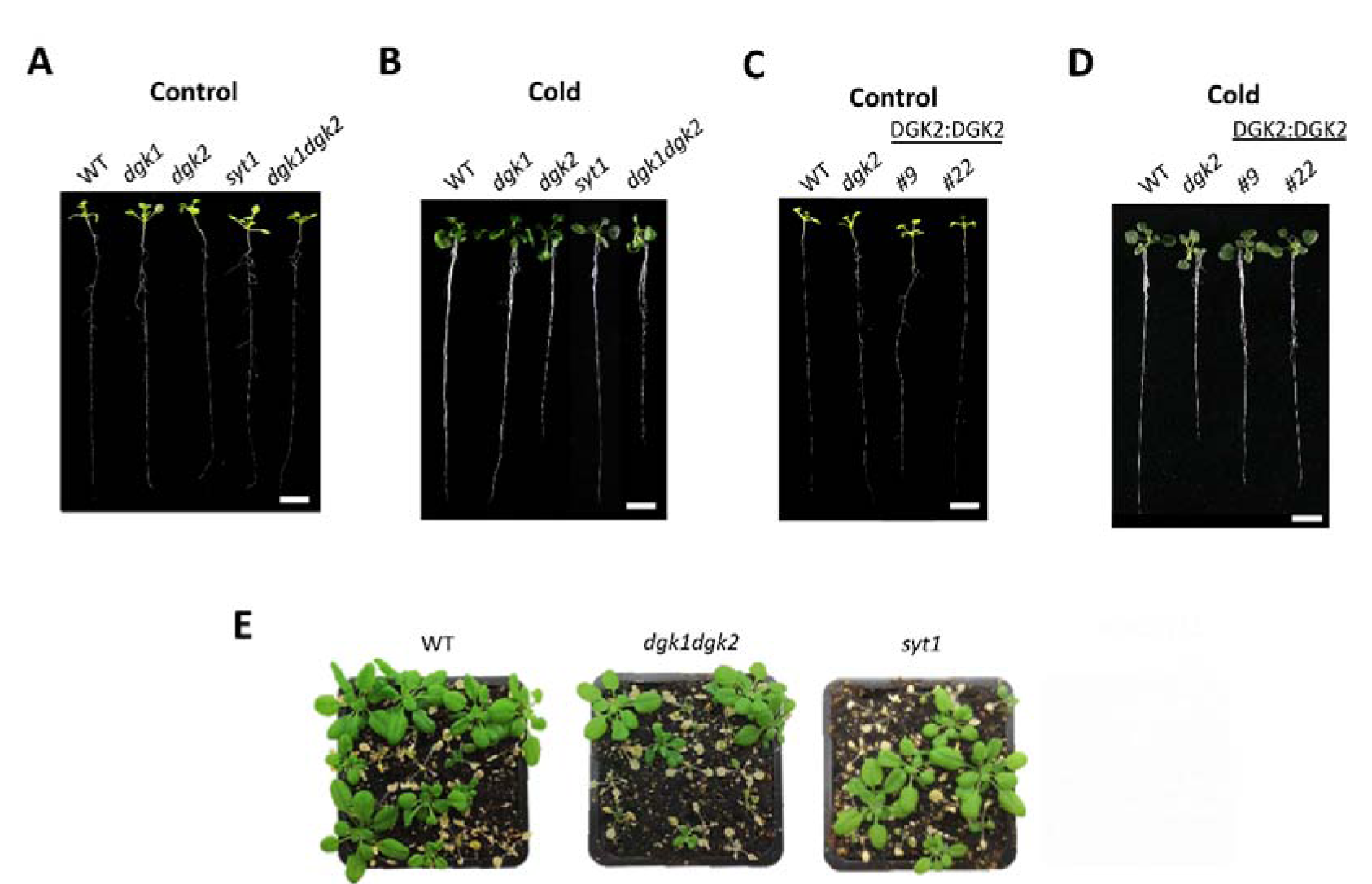
Representative seedlings from the data in Figure 7C to 7G. **A and C)** 2-week-old seedlings grown in ½ MS medium and grown vertically in control conditions (Figure 7C and 7E). Scale bar = 1 cm. **B and D)** 4-week-old seedlings grown in ½ MS medium and grown vertically 3 days in control conditions and then moved at 10°C with the same photoperiod (Figure 7D and 7F). Scale bar = 1 cm. **E)** Surviving plants of acclimated WT, *dgk1dgk2* and *syt1* after a freezing treatment 6h at -9°C and one week of recovery under control conditions (Figure 7G).

